# Effects of stress coping styles and social defeat on zebrafish behaviour and brain transcriptomics

**DOI:** 10.1101/2025.03.28.645889

**Authors:** Pavla Hubená, Lisa Benrejdal, David Brodin, Johanna Axling, Oly Sen Sarma, Peter Bergman, Svante Winberg

## Abstract

Individuals with divergent personality traits corresponding to stress coping styles have been suggested to differ in behavioural and neural plasticity. To assess the effects of social experience on the coping ability of wild zebrafish selectively bred for boldness/shyness, we used a model of social defeat stress. Behavioural tests were performed to assess parameters such as boldness/exploration, aggressiveness, and displaced behaviour. Gene expression changes in the brain were assessed via RNA sequencing. The main results suggest a strong effect of shyness and boldness phenotype on behaviour and brain transcriptome. Fish of the shy line displayed significant behavioural changes, while the number of differentially expressed genes remained low. In contrast, fish of the bold line exhibited a small effect on behaviour and pronounced changes in brain gene expression. This study highlights the importance of boldness phenotype and its influence on the response to social challenges at the behavioural and transcriptomic levels.

## Introduction

Environmental instability increases the allostatic load on the organisms, necessitating greater behavioural and physiological adaptation. Behavioural flexibility refers to the extent to which the behaviour is influenced by environmental factors, and it is crucial for successful survival^1^. Neural plasticity, which involves biochemical changes and structural reorganisation, allows animals to operate beyond their typical behavioural phenotype^2^. Consequently, individuals in fluctuating environments are likely to benefit from the ability to quickly adapt, exploit new conditions, and experience minimal adverse effects from such adaptations^3^. Individuals confront these challenges with their correlated physiological and behavioural traits, which define their “stress coping style” and predispose them to varying degrees of resilience or vulnerability in specific situations^4–6^. While current research has concentrated on the adaptability of stress coping styles in non-social contexts, such as responses to novel environments, there is a notable lack of understanding regarding how individuals with different stress coping styles adapt to negative social circumstances. Following victories or defeats, transient changes in behaviour and brain gene expression occur alongside shifts in an individual’s social rank, reflecting biochemical switching processes^7,8^. There is even less information available about the implications of such social experiences on the brain transcriptomics of individuals with varying stress coping styles, despite evidence linking defects in neural plasticity to the onset of neurodegenerative disorders^9^.

The correlated suit of behavioural traits and physiological response to stress, which tends to remain consistent in an organism over time, is referred to as “stress coping style”. The “proactive” coping style is characterised by relatively higher levels of aggression, impulsivity, boldness or risk-taking, reliance on past experiences (feed-forward control), novelty-seeking, and significant frustration behaviour.^1,10^. This proactive phenotype is primarily distinguished from the “reactive” coping style at the physiological level by a lower cortisol response to stress (indicating reduced reactivity of the hypothalamic-pituitary-interrenal (HPI) axis; hypothalamo-pituitary-adrenal (HPA) axis in mammals) and elevated plasma catecholamine levels, suggesting increased sympathetic reactivity^10–13^. Further physiological differences have been noted among various stress coping strategies, including variations in antioxidant capacity, immunological markers, brain monoaminergic activity, and indicators of neural plasticity^10,14,15^. In contrast, the “reactive” coping style demonstrates an opposing set of behavioural and physiological responses compared to the proactive style. The type and intensity of behavioural reactions contribute to individual’s resilience or vulnerability to environmental stressors. For example, proactive individuals tend to resume feeding in novel environments more quickly thanks to their stress resilience, but struggle when they cannot depend on their memory to locate food^6^. As such, proactive fish seem to thrive in stable environments that allow for routine formation, while reactive fish may excel in variable, unpredictable settings^4,6^. This difference may represent an indicator of environmentally stimulated behavioural flexibility. However, research into how behavioural flexibility is influenced by social factors has been limited to just one study^16^. The shift of the stress coping style to social challenges and its potential impact on brain transcriptomics remains an area that requires further investigation.

Social rank in dominance hierarchies, which determines the access to resources, is the result of social contests^17^. These contests are resolved through a specific set of ritualised agonistic behaviours unique to each species. In the case of zebrafish, encounters involve a variety of distinct agonistic behaviours, including displays, circling, striking, biting, chasing, retreating, fleeing, and freezing^18^. The contest ends with the establishment of a winner and a loser, with repeated interactions leading to the formation of a dominant/subordinate relationship. The outcomes of these contests have physiological effects on fish, influencing testosterone levels, HPI axis reactivity, vasotocin release, brain monoamine activation, and neural proliferation factors^7,19–25^. These physiological differences underlie the behavioural change, with winners more likely to succeed in future contests—an effect commonly referred to as the ’winner effect’. This can lead to aggressive behaviours where winners may initiate conflicts or ’bully’ the losers through biting and chasing^18,24,26^. Conversely, losers tend to exhibit a ’loser effect’, which is characterised by reduced aggressiveness, the display of submissive behaviours, and overall behavioural inhibition^18,26^. Prior social experience is not only projected to further social performance, but there is also evidence that socially dominant chickadees (*Poecile gambeli*), meadow voles (*Microtus pennsylvanicus*) and mice (*Mus musculus*) tend to outperform their subordinate counterparts in non-social tasks like caching and spatial learning^27–29^. In fish, males of African cichlid fish (*Astatotilapia burtoni*) that ascended in the social rank have been observed to alter their behaviour in non-social contexts, becoming more cautious around novel objects and hesitant to enter reward areas^23^. Mismatched or incomplete adaptation of the behaviour relative to one’s social rank within the dominance hierarchy could potentially lead to an increased number of aggressive encounters and subsequent injuries^26,30^.

Selection for boldness and social interaction has consistently been shown to affect behavioural profiles^31^ and brain gene expression^18,32^ in zebrafish, as well as in other species. Boldness has been identified as a stable behavioural trait that can predict the stress coping style in fish^33^. Nevertheless, our understanding of the interplay between this behavioural trait and social environment in shaping individual phenotypes remains insufficient.

The current study aimed to investigate the modulation of the stress coping styles in response to varied social experiences. We aimed to estimate (i) the stability of the social behaviours (aggression, displaced behaviour) between bold and shy fish before and after social interactions; (ii) the differences in non-social behaviour (boldness/exploration, activity) displayed by bold and shy fish following these social experiences; (iii) the impact of the selective breeding and social experiences on gene expression in the brain. Zebrafish, which demonstrated specific stress coping strategies, were sourced from a selection programme based on bold and shy phenotypes, connected to different stress responses^34^. The bold and shy fish were then allowed to establish a dominance hierarchy through repeated dyadic contests, ensuring that equal numbers of bold and shy fish experienced both victories and defeats. Their behavioural profiles and gene expression were subsequently analysed. Gene expression analysis was conducted via bulk RNA sequencing on brain tissue samples, enabling comprehensive transcriptomic profiling across different fish lines following social exposure.

## Materials and Methods

The current study used zebrafish males originating from artificial selection programme focusing on differences in boldness and shyness as assessed by the NTD test. Description of the selection programme is provided in Supplementary files (**SF-1)**. A total of 40 adult male zebrafish from the F2 generation were used, comprising 20 from the bold line and 20 from the shy line. The fish had no prior experience of behavioural testing or any other procedure. The fish from the selection programme were housed in Aquaneering zebrafish racks, in 6-9 L tanks. The temperature was 27.2 ± 0.3 °C, pH 8.5 and the NH3/NH4^+^ was 0 in the housing racks. During the experiment, the fish were housed in 6 L tanks (29 × 20 × 15 cm) containing a filter (Eheim GmbH & Co, Germany) and heater tempering the water to 27.9 ± 0.5 °C (mean ± SD). The NH3/NH4^+^ reached 0.7 ± 0.3 mg/L and the pH was 8.5 throughout the whole experiment. Half of the tank volume (3 L) was exchanged once a day for aged, heated (28 °C), and aerated water. The photoperiod was maintained at 14 hours of light and 10 hours of darkness. The fish were fed twice a day with standard feed for zebrafish in laboratory conditions (Zebrafeed 400-600 mm, Sparos, Portugal) and rotifers (*Brachionus* spp.).

### Experimental design

The timeline of the experiment was illustrated in the **Fig. 1**. Four days prior to the beginning of the experiment, fish were anaesthetized in the immersion of 200 mg/L of benzocaine and were immediately transferred to experimental tanks, each tank holding a pair of fish, size matched based on their body weight (10% difference at maximum). The pairs consisted of familiar fish originating from the same selected line group (bold or shy). The dorsal or the ventral side of the caudal fin of the fish was clipped to allow visual determination of individuals in each pair. The individuals in an experimental tank were divided by a liftable opaque barrier. The fish were allowed to acclimate to these conditions for three days. Following acclimation, the isolated fish were first allowed to fight with its mirror image in the mirror test (MT) in order to monitor differences in aggression and displaced behaviour prior to the social experience. Displaced behaviour is tested more in rodents (e.g., grooming) than in fish but has been found to increase in response to stressful situation as a result of activation of the limbic system^35,36^. During the MT, a mirror was slowly inserted into the compartment of individual fish and their reaction was recorded for 10 minutes using a GigE camera. Following the MT, the partition separating fish in pairs was removed and the fish were allowed to interact with their size-matched opponent. In order to establish repeated winning or losing experience, the fish were allowed to interact for three days, 5 hours a day. The outcome of the contests was visually assessed based on the subordinate behaviour (“tail drop”), initiation of fights by the winner and occupation of the central part of the tank by the winning fish. In all pairs, a clear dominant and subordinate status was established after three days of social contest. After 3 days of repeated dyadic interaction, the fish were tested in the zebrafish Multivariate Concentric Square Field (zMCSF) test to monitor non-social behaviours. After this behavioural screening, the fish were returned to their experimental tanks and subjected to three more days of dyadic interaction with their size matched opponent, 5 hours a day, in order to re-establish the social experience. Following the dyadic contests, the fish were tested again in the MT in order to monitor effects of social experience on aggression and displaced behaviour. After this final mirror test, the fish were immediately sacrificed using 100 mg/L of benzocaine bath and their brain was sampled within 3 minutes from euthanasia.

**Fig. 1:**
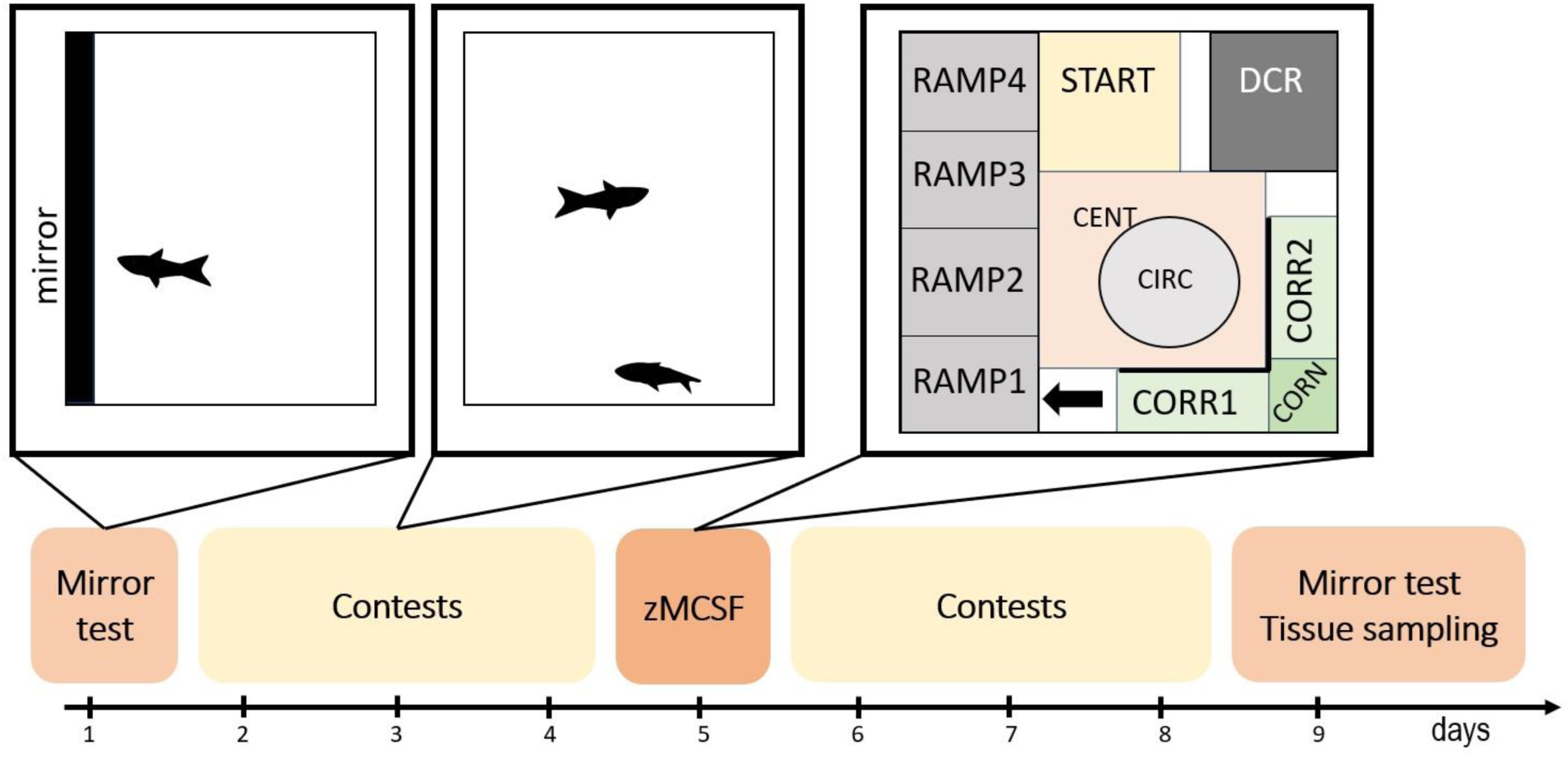
The experimental design of the experiment including the mirror test assays, dyadic contests and the zebrafish Multivariate Concentric Square-Field (zMCSF) test. The individual tests are visualized above the timeline. CENT = central zone, CIRC = circular zone, CORN = corner zone, CORR1 = corridor 1, CORR2 = corridor 2, DCR = dark corner with a roof, RAMP (1-4) = inclined ramp with RAMP4 having the shallowest water depth, START = start zone. Unmarked zones belong to the REST zone. Black arrow indicates entry to RAMP (1-4) zones.

### zMCSF behavioural assay

Square opaque arena (30 × 30 × 25.8 cm) was filled with 8 L of aged, aerated, and pre-heated copper-free Uppsala municipal tap water (27.5 ± 0.3 °C (mean ± SD)). Black plexiglass walls were placed at a 90° angle around a corner to create a corridor (zones CORR1, CORN, CORR2). A second corner was sheltered by black plexiglass roof, which created a dark roofed corner (DCR). An inclined transparent ramp was placed along the opposite wall (zones RAMP1, RAMP2, RAMP3, RAMP4) and provided gradually reduced water depth. In order to prevent the preference of fish to specific zones based on external cues, the experiment was carried out using 4 arenas at the same time and each arena was rotated in 90° angle from other arenas. These arenas were placed on top of an infrared backlights (Noldus, Wageningen, the Netherlands) and the trial was recorded by infrared camera (Basler GigE camera) connected to the tracking system Ethovision XT 16 (Noldus, Wageningen, the Netherlands). The whole apparatus was enclosed in a separate behavioural room to prevent any disturbances. In this test, movement through the corridors is interpreted as locomotor activity and the movement through ramps and the open field area is translated as boldness/exploration, as described by Vossen et al.^37^.

Two experimental tanks containing a random pair were placed into the behavioural room for five minutes for acclimation. Each individual was then netted from their compartment and placed into the START zone in their arena, where they were allowed to freely explore the arena for 30 minutes. Their behaviour was recorded by the above-placed camera and tracked in Ethovision XT 16. After the trial, the fish were transferred from the arena back into their original compartment and the tank was placed back into its original location. The temperature in the arena was controlled before and after the trial.

### RNA isolation and RNA sequencing

A vertical section between the cerebellum and the optic tecta was performed to divide the brain into forebrain and midbrain (FBMB) and hindbrain (HB). All samples were stored for 24 hours in 500 μL of RNA later in 4 °C after which they were relocated to -20 °C until the RNA isolation. A total of 12 pairs (n = 24 fish) was included in the transcriptomic analysis. The brain tissues were then transferred from the RNA later to 300 μL of RLT lysis buffer and homogenised using the TissueRuptor II (Qiagen, Inc.). RNA isolation was performed using the RNeasy Fibrous Tissue Mini Kit (Qiagen, Inc.) according to the manufacturer’s protocol. Total RNA was subjected to quality control with Agilent Tapestation according to the manufacturer’s instructions. To construct libraries suitable for Illumina sequencing the Illumina Stranded mRNA Prep, Ligation preparation protocol was used which includes mRNA isolation, cDNA synthesis, ligation of anchors and amplification and indexing of the libraries. The yield and quality of the amplified libraries was analysed using Qubit by Thermo Fisher and the Agilent Tapestation. The indexed cDNA libraries were finally normalised and combined, and the pool was sequenced on two lanes on the Illumina Novaseq 6000 plus 10 B flowcell, 2x 150 bp, paired end mode generating a total of X reads.

### Ethical statement

The study was authorized by the Uppsala Animal Ethical Committee (permit 5.218-13393/2023). All procedures followed the guidelines of the Swedish Legislation on Animal Experimentation (Animal Welfare Act SFS 2018:1192) and the European Union Directive on the Protection of Animals Used for Scientific Purposes (Directive 2010/63/EU).

### Data analysis

‘Line’ category contained fish with different behavioural phenotypes, bold (‘B’) or shy (‘S’). ‘Rank’ was including divergent social experiences the fish received, losing (‘L’), or winning (‘W’). ‘Zone’ category contained 12 zones present in the zMCSF trial.

The ethological software BORIS^38^ was used to score aggressive and displaced behaviour from the mirror test assay. The aggression was estimated based on the number, total duration, latency to first attack, and average duration of attacks. The average duration of attack was calculated by division of the total duration of attack by the number of aggressive interactions. For aggressive behaviour to be recorded in zebrafish, they must have made physical contact with the mirror, which indicates overt aggression. Displaced behaviour was scored as the number, total and average duration, and latency to the first expression of the behaviour. Specifically, the fish exhibited a sudden compulsion to search for and bite at non-existent food, either from the bottom or the top of the tank. This was classified as displaced behaviour because food was not made available to the fish during the contest, and their reaction appeared as an abrupt behavioural shift that was inappropriate to the context of the threat posed by the mirror image.

Ethovision XT 16 software was used for ethological evaluation of the recorded videos from the zMCSF test. Total distance moved (cm), average velocity (cm/s), and the immobility duration (s) were extracted for each arena. Total activity was calculated as the sum of all zone entries and was evaluated only during the whole-arena analysis. For zone-specific analysis, total distance moved (cm), average velocity (cm/s), the number of visits to each zone, latency to first entry to the zone (s) and cumulative duration in each zone (s) were extracted from the Ethovision software. Duration of stay in each zone was calculated by dividing the cumulative duration of stay in zone by the duration of stay in the whole arena. Visit duration in each zone was calculated as the cumulative duration in each zone divided by the number of entries to the zone and the total value was divided by the total duration of stay in the arena. The number of entries to zone (%) was calculated by subtraction of the number of entries to zone by the total activity and the result was divided by the value of 100.

The transcriptomic data were analysed as following: Basecalling and demultiplexing were performed using Illumina bcl2fastq (v2.20.0). Sequence data quality was assessed using FastQC (v0.11.8). Reads were aligned to the Ensembl Danio rerio GRCz11 reference genome using STAR (v2.7.9a).

Due to technical issues, four pairs of the fish (n = 8) had to be excluded from the final analysis. The number of fish per each selected line and social experience was equal (n = 8 per each group; n = 32 in total).

### Statistical analysis

The behavioural analysis was performed in R version 4.3.0 ^39^. The packages ‘bestNormalize’ ^40^, ‘ggplot2’^41^, ‘emmeans’^42^, ‘lme4’^43^, ‘car’^44^, and ‘MASS’^45^ were used to analyse the behavioural dataset.

The aggressive and displaced behaviour was evaluated before the social experience for establishing differences between the behavioural phenotypes, i.e., the selected lines. Therefore, the Generalized Linear Mixed Effect Models (GLMM) with Poisson distribution and Line as fixed effect and individual as random effect were used to analyse the number of aggressive and displaced behaviours before the social experience. The latency to first attack or displaced behaviour, the total duration of attack or displaced behaviour, and the mean duration of attack or displaced behaviour were evaluated using one-way ANOVA with Line as explanatory variable. The percentage change was used to analyse the effect of social experience, i.e., the change of behaviour from its original value. The percentage change of all aggressive and displaced behaviours was evaluated using two- way ANOVA with Line and Rank as explanatory variables. The zMCSF behavioural assay was analysed as a behaviour in the whole arena and in specific zones. The total distance moved, average velocity, and the immobility duration in the whole arena in the zMCSF test was analysed using two-way ANOVA with Line and Rank as independent variables. The total activity in the zMCSF arena and the number of visits per zone were analysed using the Generalised Linear Mixed Model (GLMM) with Negative Binomial distribution, as this model yielded better fit (according to AIC) than the Poisson distribution. The total distance moved, average velocity, the duration in each zone, visit duration, number of entries to zone (%), and latency to first entry were evaluated using the Linear Mixed Effect Models (LMM). Line and Rank were used as fixed factors, and individual was applied as a random variable. Zone was added as fixed factors for all zone-specific parameters in the zMCSF behavioural assay.

Normal distribution was assessed using the Shapiro-Wilk test of normality. The transformation method was recommended by the ‘bestNormalize’ package. Each parameter was transformed according to recommended transformation if the normal distribution was not reached (for details see **SF-2**). emmeans function (‘emmeans’ package) with Bonferroni correction was used to observe pairwise comparison of factors in GLMMs, LMMs, and one- and two-way ANOVAs, and Bonferroni correction was used to adjust the p-value. Significant differences were evaluated based on α = 0.05. The weight of the fish was not significantly different between groups at the beginning of the experiment and was omitted from the analysis.

Counts for each gene were estimated using featureCounts (v1.5.1). The Bioconductor package DESeq2 (v.1.42.0) was used for sample group comparisons, generating log2 fold changes, Wald test p-values and p-values adjusted for multiple testing (Benjamini-Hochberg method).

## Results

The objective of the experiment was to investigate changes in behaviour resulting from various social experiences. To identify behavioural differences in aggression, displaced behaviour, boldness, and activity, a mirror test (MT) and the zebrafish Multivariate Concentric Square Field (zMCSF) test were employed (**Fig. 1**). Additionally, the impact of selection for boldness and social experiences on the brain transcriptome was assessed through RNA sequencing of brain samples obtained post-behavioural testing.

Generally, the stress coping style (see selection program in **SF-1**) produced more differences than the recent social experiences. This distinction was evident in the behavioural analyses (see **Table 1**, **Table 2, SF-2**) and the brain transcriptomics (**SF-3**).

**Table 1:** The initial differences between the selected lines in the aggressive and displaced behaviours (Trial 1) and the differences between groups in the percentage change of the parameters from the first to second trial indicating the effect of social experience (Trial 1 - 2). B = bold selected line; BL = bold loser; BW = bold winner; NS = not significant; S = shy selected line; SL = shy loser; SW = shy winner. ***p < 0.001; **p = (0.01; 0.001); *p = (0.5; 0.1)

|  | **Trial 1** | **Trial 1 - 2** |
| --- | --- | --- |
| **Aggression** |  |  |
| Number of attacks | B > S** | BW < SW* |
| Latency to attack | NS | BL < SL* |
| Total duration of attack | B > S* | NS |
| Average duration of attack | B > S* | NS |
| **Displaced behaviour** |  |  |
| Number of displaced behaviours | NS | NS |
| Latency to displaced behaviour | NS | NS |
| Total duration of displacement | NS | NS |
| Average duration of displacement | NS | NS |

**Table 2:** A summary table over zMCSF parameters for which there were significant differences between bold and shy winners and losers. AVEL = average velocity (cm/s); DUR = duration in zone (s); LAT = latency to first entry (s); NVIS = number of visits to zone; NVIS (%) = number of visits to zone in percentage; TDIS = total distance moved (cm), VDUR = visit duration (s).

| Functional categories | Parameters | Selected lines | Social rank |
| --- | --- | --- | --- |
| Boldness/exploration | TDIS RAMP2 | BL > SL* |  |
| Locomotor activity | TDIS RAMP3 | BL > SL* |  |
|  | TDIS RAMP4 | BW > SW*** |  |
|  | AVEL CENT | BW > SW* |  |
|  | AVEL CIRC |  | SL > SW* |
|  | DUR RAMP2 | BL > SL* |  |
|  | DUR RAMP3 | BL > SL* |  |
|  | DUR RAMP4 | BL > SL* |  |
|  |  | BW > SW*** |  |
|  | VDUR RAMP4 | BW > SW** |  |
|  | NVIS RAMP2 | BL > SL* |  |
|  | NVIS RAMP3 | BL > SL* |  |
|  | NVIS RAMP4 | BL > SL* |  |
|  |  | BW > SW*** |  |
|  | NVIS (%) RAMP2 | BL > SL* |  |
|  | NVIS (%) RAMP3 | BL > SL* |  |
|  | NVIS (%) RAMP4 | BW > SW*** |  |
|  | TDIS CORR2 | BW > SW* |  |
|  | TDIS CORN | BW > SW* |  |
|  | DUR CORR2 | BW > SW* |  |
|  | DUR CORN | BW > SW** |  |
|  | NVIS CORR2 | BW > SW* | SL > SW* |
|  | NVIS (%) CORR2 | BW > SW** | SL > SW* |
|  | NVIS (%) CORN | BW > SW* |  |
|  | LAT CORR1 | BW < SW* |  |
|  | LAT CORR2 | BW < SW* |  |
|  | LAT CORN | BW < SW* | BL > BW* |

### Aggression and displaced behaviour

There were significant differences in aggression and displaced behaviour before any social experience was induced indicating a significant effect of the selected line (**Table 1**). Specifically, bold fish performed significantly higher number of attacks (GLMM: z = 2.783, p = 0.005), higher duration of attacks (ANOVA: t32, 30 = 2.712, p = 0.011), and higher average duration of attack (ANOVA: t32, 30 = 2.453, p = 0.02) than shy fish. The social experience was not considered in this analysis as the fish had not been exposed to the social experience at that time and were isolated for three days prior to testing.

The percentage change reflects the variation in behaviour that occurred during the social experience. There were no significant changes from the baseline value (p > 0.05), but some groups differed in their percentage change. The percentage change of the number of attacks of bold winners was significantly lower than that of shy winners (ANOVA: t32, 28 = -2.632, p = 0.014). The percentage change of the latency to the first attack was significantly lower in bold losers than in shy losers (ANOVA: t32, 28 = -2.277, p = 0.031). The magnitude of the percentage change indicates how plastically the fish reacted to altered social experience, indicating that both shy winners and losers were more flexible in their behaviour than their bold counterparts.

### Zebrafish Multivariate Concentric Square Field test

When specific zones were not accounted for, the total distance moved, average velocity, total activity and immobility was not significantly different between any selected line or social experience. Significant differences were found only when the zone factor was added to the calculation (see individual zones in **Table 2**). There were no differences in any parameter in START, DCR, REST, and RAMP1 zones. However, there were differences in CORR1/CORN/CORR2, RAMP2/RAMP3/RAMP4, and CENT/CIRC zones. For the full table see (**SF-2**).

Selected lines and social rank significantly affected the behaviour of fish in the CORR1/CORN/CORR2 zones. Bold winners entered CORR1 faster than shy winners (LMM: t366, 288 = 2.276, p = 0.024). Bold winners also entered CORN zone faster (LMM: t366, 306 = 2.480, p = 0.014), visited this zone more if compared in percentages (LMM: t366, 306 = 2.480, p = 0.014), spent a longer duration of time in this zone (LMM: t366, 260 = 2.651, p = 0.009), and travelled longer distances in this zone (LMM: t366, 171 = 2.145, p = 0.033) than shy winners. Similarly, bold winners entered CORR2 zone faster (LMM: t366, 288 = 2.075, p = 0.039), visited this zone more (Negative Binomial GLMM: z = 2.331, p = 0.020; in percentages: LMM: t366, 306 = 2.904, p = 0.004), stayed there longer duration of time (LMM: t366, 260 = 2.272, p = 0.024), and travelled longer distances in this zone (LMM: t366, 171 = 2.533, p = 0.012) than shy winners. Social experience also significantly diversified the behaviour in CORN and CORR2 zones. Specifically, bold losers entered the CORN zone later than bold winners (LMM: t366, 290 = 2.151, p = 0.032) and shy losers visited the CORR2 zone more than shy winners (Negative Binomial GLMM: z = 2.367, p = 0.018; in percentages: LMM: t366,308 = 2.162, p = 0.031).

There were significant differences in behaviour in RAMP (2-4) zones and CIRC/CENT zones between the selected lines, but no effect of social experiences. Specifically, bold losers visited RAMP2 more (Negative Binomial GLMM: z = 2.236, p = 0.025; in percentages: LMM: t366, 307 = 2.315, p = 0.021), spent there longer duration of time (LMM: t366, 271 = 2.356, p = 0.019) and travelled longer distances there (LMM: t366, 190 = 2.447, p = 0.015) than shy losers. These differences were also significant in RAMP3 zone, as bold losers visited RAMP3 more (Negative Binomial GLMM: z = 2.480, p = 0.013; in percentages: LMM: t366, 306 = 2.090, p = 0.037), spent a longer duration of time there (LMM: t366, 265 = 2.345, p = 0.020) and travelled longer distances in that zone (LMM: t366, 180 = 2.575, p = 0.011) than shy losers. Both bold winners and losers visited the RAMP4 zone more (Negative Binomial GLMM: W: z = 3.945, p <0.001; W (in %): LMM: t366, 307 = 4.029, p <0.001; L: z = 2.176, p = 0.030) and spent longer duration of time in this zone (LMM: W: t366, 268 = 4.055, p <0.001; L: t366, 265 = 2.158, p = 0.032) than shy winners and losers, respectively. However, only bold winners spent significantly longer duration per each visit in this zone (LMM: t366, 250 = 2.634, p = 0.009) and travelled longer distances through this zone (LMM: t366, 182 = 3.594, p = 0.004) than shy winners. This indicates that shy winners exhibited a comparable behaviour to bold winners up to the RAMP3 but avoided the RAMP4 zone more.

Zebrafish also differed significantly in average velocity in the CENT/CIRC zone, which was based on selected line and social experience. Bold winners swam significantly faster through the CENT zone than shy winners (LMM: t366, 111 = 2.492, p = 0.014), but the average velocity only approached significance level between bold and shy losers (LMM: t366, 111 = 1.967, p = 0.052). There was a difference between the selected lines depending on the social experience: shy losers moved significantly faster through CIRC zone than shy winners (LMM: t366, 126 = 2.001, p = 0.048). Therefore, the social defeat seems to have caused increase in swimming speed through the CIRC zone in shy, but not in bold fish.

### Transcriptional changes in the brain related to stress coping styles and social interactions

Principal Component Analysis was used to evaluate the general differences in brain transcriptomic profiles between the selected lines and social experiences (**Fig. 2**). The PC1 was explained by the difference between the sampled areas of the brain (FBMB, HB) at 58.5 % variance. PC2 of the PCA was explained by the bold and shy stress coping style at 5.5 % variance. No significant distinction between winners and losers could be estimated based on the PCA. Thus, the largest variability in the PCA-analysis could be explained by the brain regions, followed by the stress- coping styles and social experience.

**Fig. 2:**
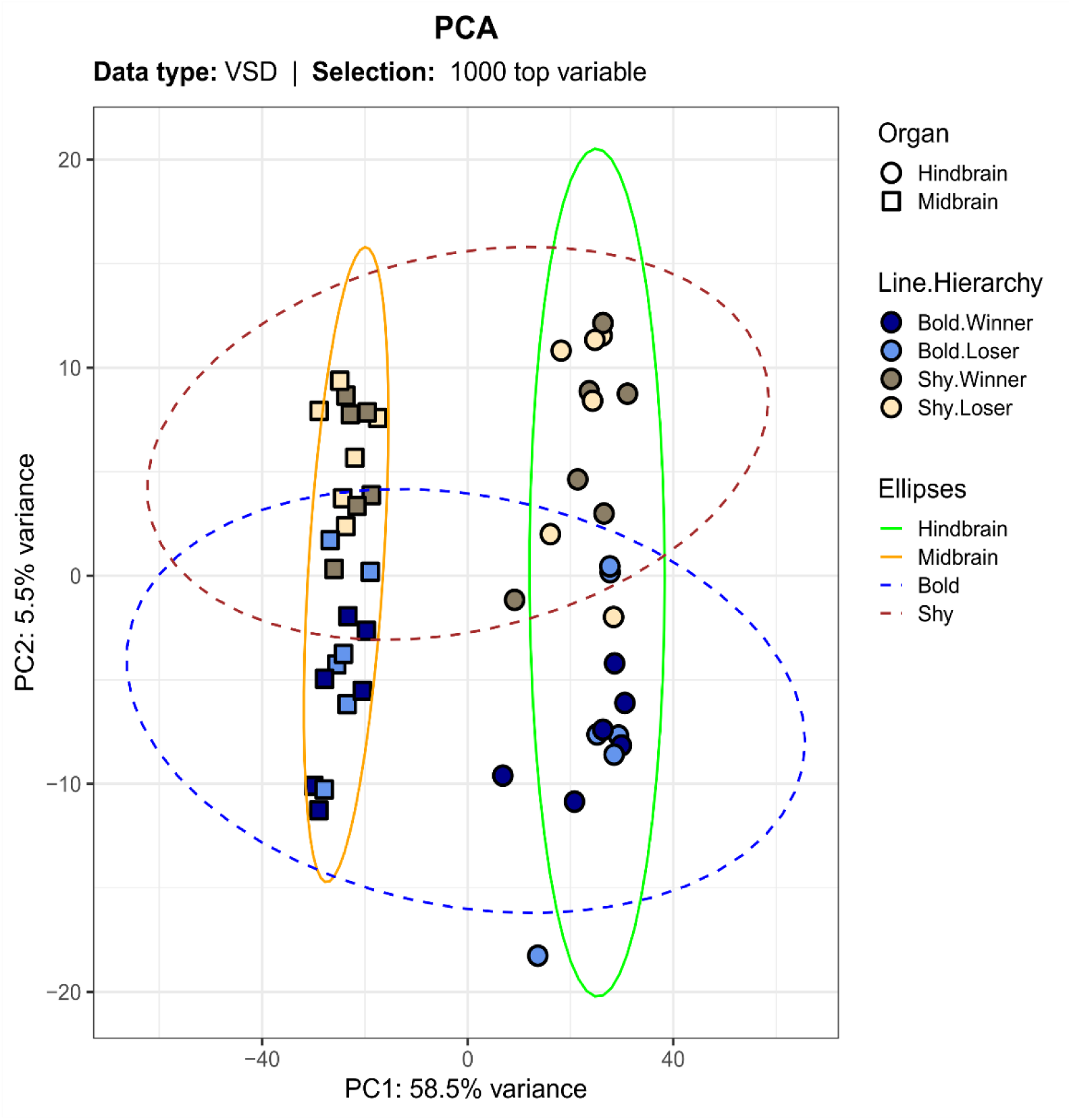
The Principal Component Analysis (PCA) including 1000 top variable genes in the Forebrain/Midbrain and Hindbrain of the groups defined by line (shy/bold) and social experience (loser/winner). Four clusters described by brain region and fish line are highlighted by ellipses.

The differences observed in the PCA were reflected in the number of differentially expressed genes (DEGs) (**Fig. 3**). There were marked differences based on selected line within particular brain regions (see **SF 3**). In hindbrain, the shy losers had downregulated the immediate early response genes (*ier2b*), neuronal PAS domain protein 4a (*npas4a*), and nuclear receptor subfamily 4, group A, member 1 (*nr4a1*) involved in neuronal development and plasticity in comparison to bold losers. Additionally, several genes linked to possible oncogenesis (*ccnb1, plk1, pim1, pttg1, spry4*) or senescence (*pim1, gapdh, ldhbb*) were downregulated, except for the *ldhbb* gene, which was upregulated (**Fig. 4A**). In forebrain/midbrain, the losers of shy vs bold line manifested differences in a particular gene: WD repeat domain 45 (*wdr45*) was upregulated in shy losers compared to bold losers (**Fig. 4B**). The shy winners showed upregulation in the forebrain/midbrain region in genes contributing to the neuronal development, plasticity and memory formation, such as the proline dehydrogenase 1a (*prodha*), toll-like receptor 7 (*tlr7*), and vimentin-like *(viml*) compared to bold winners. The corticotropin releasing hormone receptor 1 (*crhr1*) involved in the HPI axis of fish was upregulated in shy winners compared to bold. DEGs of other significant processes in the forebrain/midbrain were downregulated in shy winners compared to bold: genes involved in iron metabolism (*fthl29, slc11a2, tfr1a)*, and pyroptosis (*gsdmea, rflnb*). All were downregulated in shy vs bold winners, apart from the *rflnb* gene (**Fig. 4C**). Despite the significant change in gene expression, some of the genes (*nr4a1, spry4, ldhbb, pim1, wdr45, tlr7, rflnb, prodha, crhr1, gsdmea, slc11a2*) did not reach the pre-defined cutoff of 2-fold change.

**Fig. 3:**
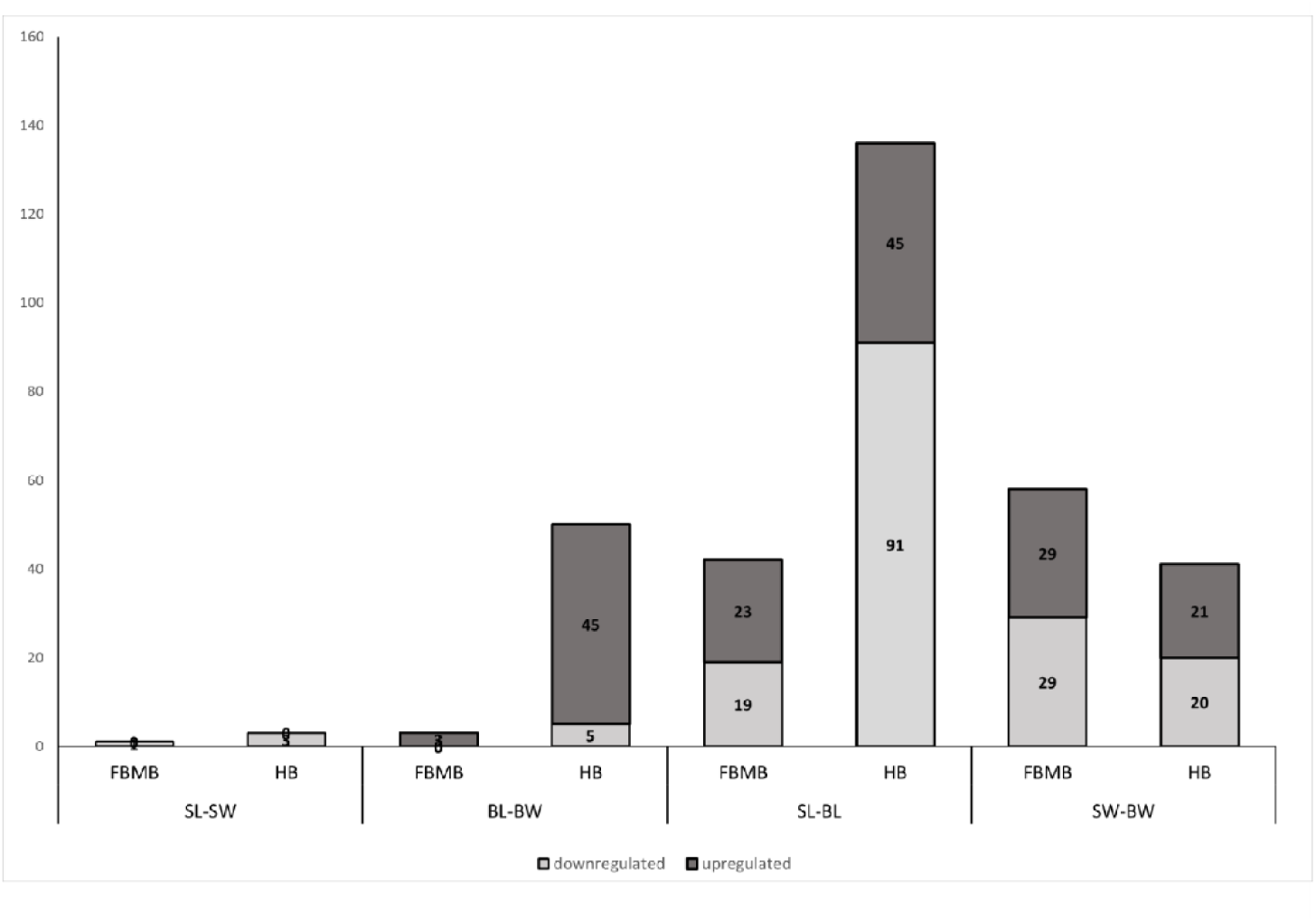
Number of differentially expressed genes in Forebrain & Midbrain (FBMB), Hindbrain (HB) of groups with divergent behavioural phenotypes and social experience. SL-SW = Shy Loser vs Shy Winner; BL-BW = Bold Loser vs Bold Winner; SL-BL = Shy Loser vs Bold Loser; SW- BW = Shy Winner vs Bold Winner.

**Fig. 4:**
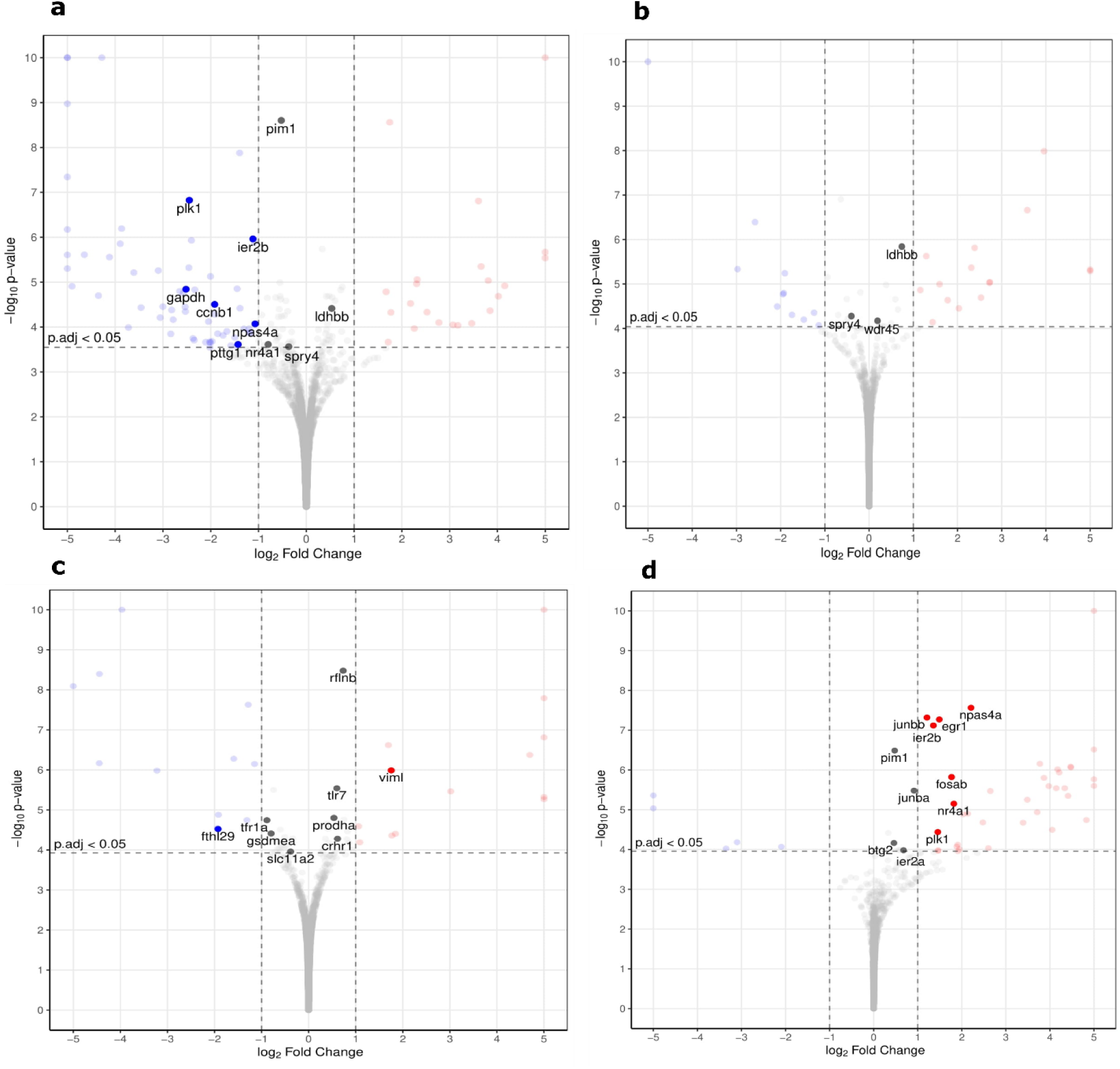
Volcano plots representing the differentially expressed genes in **a.** HB shy losers vs bold losers; **b.** FBMB shy losers vs bold losers; **c.** MB shy winners vs bold winners; **d.** FBMB bold losers vs bold winners. Red dots represent upregulated transcripts, and blue dots represent downregulated transcripts in the given group (*p*-value ≤ 0.05 and log2 Fold Change > 1).

The bold and shy fish responded differently to social challenges. There were few transcriptomic changes between winners and the losers in the shy line in any of the brain regions. Specifically, the shy fish differed very little in their transcriptomic profiles, which was intriguing. Bold fish, however, responded significantly to social experiences. The socially regulated genes in the hindbrain in the bold line (loser vs winner) included upregulation of the B-cell translocation gene 2 (*btg2*), early growth response genes (*egr1*), FBJ osteosarcoma oncogene (*fosab*), immediate early response genes (*ier2a, ier2b*), jun B proto-oncogene (*junba, junbb*), neuronal PAS domain protein 4a (*npas4a*), and nuclear receptor subfamily 4, group A, member 1 (*nr4a1*), which are involved in neuronal development and plasticity (*btg2, egr1, junba, junbb, npas4a, nr4a1*). Specific genes, such as *pim1* and *plk1* were upregulated, which are potentially linked to oncogenesis (**Fig. 4D**). The pre-defined cutoff value of 2-fold change was not reached by *pim1, junba, btg2* and *ier2a*, even though their expression was significantly altered. In the forebrain/midbrain region, only a single gene was differentially expressed between shy winners and losers, and only 3 DEGs were detected between the bold winners and losers, which precludes solid conclusions based on these changes. Combined, the effects of the social experience were most pronounced in bold fish and involved genes related to neural development, plasticity, and potentially oncogenesis.

## Discussion

This study demonstrates that zebrafish selected for distinct behavioural phenotypes exhibit differing responses to social interactions. Furthermore, these variations in behaviour are associated with differences in the transcriptomic profiles of the forebrain/midbrain and hindbrain in bold and shy fish. However, our findings indicate that in zebrafish, the intrinsic behavioural profile, or stress coping style, is a more significant influence on behaviour and brain gene expression than the experience of social interaction.

### Stress coping styles

Our results demonstrate significant behavioural and transcriptomic differences between male zebrafish selected for contrasting boldness phenotype. Fish selected for boldness or shyness were tested first for aggression and displaced behaviour, which allows clear distinction between bold and shy fish without considering social rank (winner and loser). Consequently, the other behavioural parameters were tested after repeated social experience and, hence, the social experience (winner, loser) was considered in the analysis. In other words, the stress coping styles are observed as differences between bold and shy winners as well as between bold and shy losers. The bold individuals exhibited increased aggression (3 significant differences) prior to any induced social interactions. They also demonstrated significantly enhanced boldness and exploratory behaviours (16 significant differences), along with increased locomotor activity in the zMCSF arena (10 significant differences). The variation in risk-taking behaviour or boldness has been shown to differ significantly among proactive and reactive individuals in early developmental stages of zebrafish, flathead grey mullet (*Mugil cephalus*), common carp (*Cyprinus carpio*), and sea bream^33,46–49^. In this study, we show that selection for boldness has produced behavioural differences that correlate with stress coping phenotypes. Alternatively, lines may be selected based on post-stress cortisol levels or hypoxia tests ^50–52^, although research by Alfonso et al.^33^ indicates that fish exhibit greater consistency in selection for risk-taking behaviour due to its strong association with other behavioural indicators—including exploration of novel environments, locomotor activity, and competitive ability/aggression—forming a behavioural phenotype characteristic of a proactive stress coping style^46,49,50^. The selection programme in this study produced significant differences within the population, which behaviourally align with the distinctions between proactive and reactive stress coping styles^53^. These differences were distinctly observable despite varying social experiences, demonstrating consistency over time and across different situations^1^.

The selected lines are anticipated to exhibit variations in stress responses, which will subsequently be apparent in their transcriptomic profiles. A key characteristic of stress coping styles is the divergence in reactivity of the hypothalamus-pituitary-interrenal (HPI) axis ^10^. In the present study, bold winners demonstrated a significantly lower expression of the CRH1 receptor compared to shy winners, indicating a reduced responsiveness to stress via the HPI axis in this cohort. Following the activation of the HPI axis, the stress hormone cortisol is released^54,55^. Despite the significant upregulation, the pre-defined threshold of a 2-fold change was not achieved by the *crhr1* gene in winners, which may be attributable to the small sample size (n = 6) of fish per group utilised for transcriptomic analysis or influenced by other underlying factors. Notably, bold losers were indistinguishable from shy losers regarding the expression of stress markers, suggesting that recent social experiences had enduring effects on the reactivity of the HPI axis, thereby diminishing selected line differences. It has been suggested that low social rank may lead to chronic stress, as these individuals typically display behavioural inhibition and submissive postures^56,57^. The expression of the CRH1 receptor could have been increased in bold losers, likely as a result of the chronic stress they encountered, given that the CRH1 receptor is associated with anxiety ^58^.

The transcriptomic profiles of shy and bold fish are influenced by their social experiences. Bold vs shy losers had upregulated the immediate early genes involved in the mitogen activated protein kinase (MAPK) pathway (*ier2b, npas4a, nr4a1*; WikiPathways; STRING database; **Fig. 5**). These genes function as transcription factors, regulating essential cellular processes such as the cell cycle and the balance between excitatory and inhibitory signals^59–61^. Chronic social defeat has been shown to elevate the expression of these genes, as observed in studies involving mice^62^. Typically, these genes are associated with neural plasticity^62–64^. While it is generally believed that shy or reactive lines exhibit greater plasticity^6^, their reduced response may also stem from differences in the intensity of the social defeat experienced compared to their bold or proactive counterparts. Shy fish tend to be less aggressive than bold fish, suggesting that social defeat might be less detrimental for them, resulting in a milder transcriptomic response^65^. Nevertheless, all pairs displayed clear dominant/subordinate postures following dyadic contests, irrespective of their behavioural line. Conversely, shy losers exhibited an upregulation of the *wdr45* gene in the forebrain/midbrain, which is known to enhance cognitive function^66^. Even though the stress did not induce as prominent activation of the MAPK pathway in shy losers that would support the neural plasticity proposition, they had a predisposition for learning ability due to the *wdr45* gene in the forebrain/midbrain region.

**Fig. 5:**
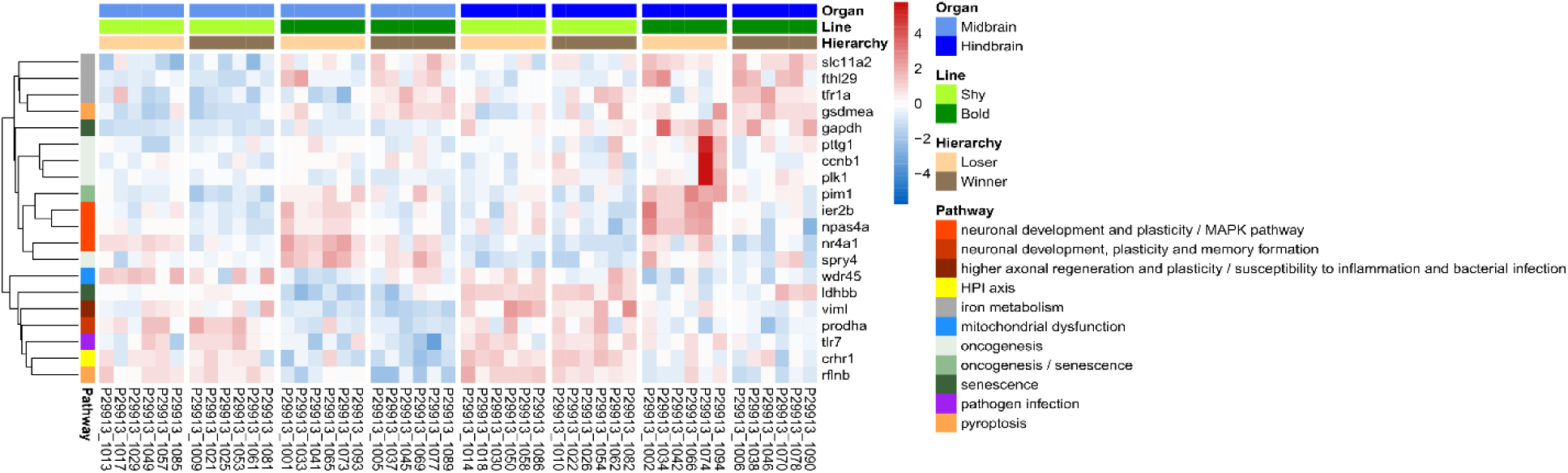
Graphical representation of biological processes and pathways associated with differentially expressed genes in the midbrain/forebrain and hindbrain between groups from different lines and social experience. The pathways were collected from literature search and based on the STRING database (https://string-db.org/).

The bold losers exhibited notable differences from shy losers in several oncogenic markers, including *ccnb1, plk1, pim1, pttg1,* and *spry4*, which were found to be upregulated. The STRING database indicated that the *plk1, pttg1* and *ccnb1* genes are co-expressed and connected to the cell cycle pathways in KEGG and WikiPathways. Current literature suggests that these markers are more closely associated with the growth of microglia and astrocytes rather than neuronal proliferation^67–70^. Consequently, bold losers are at a significantly higher risk for microglial cell proliferation compared to their shy counterparts, which demonstrate the expression of several genes linked to cellular arrest, apoptosis, and senescence, such as *pim1, gpdh, ldhbb*^71^. It is worth noting that many genes influencing the cell cycle showed only a modest increase in expression levels and were thus below the pre-established threshold of a 2-fold change. Therefore, we interpret this finding as an indication of predisposition rather than definitive causation.

The analysis of the transcriptomic profiles of bold and shy winners revealed significant differences, suggesting distinct predispositions compared to those observed between bold and shy losers. In terms of neural plasticity, shy winners exhibited upregulation of the vimentin-like gene (*viml*). This gene’s corresponding protein serves a dual role; while it stabilises the intracellular structures of intermediate filaments and enhances axonal regeneration and plasticity, it concurrently increases the risk of susceptibility to inflammation and bacterial infections^72,73^. Shy winners, therefore, appear to be predisposed to greater synaptic plasticity, but also potentially more vulnerable to pathogen infections. This is further evidenced by the overexpression of the toll-like receptor 7 (*tlr7*) gene, which has been associated with heightened levels in tissues affected by pathogen infections^74^. One of the primary mechanisms for pathogen elimination is pyroptosis, a form of cell death initiated by inflammasomes^75^. The gene expression from the current study indicates that shy winners were indeed in a higher risk of pathogen infection, as demonstrated by the expression levels of genes associated with pyroptosis (upregulation of *rflnb* and downregulation of *gdsmea*)^76^ which could also lead to neuroinflammation. Prior research has shown that chronic elevation of stress hormones correlates with increased mortality from common fungal and bacterial diseases^77^. More recently, reactive fish, classified based on post-stress cortisol levels and skin pigmentation, have been shown to be more susceptible to infections from sea lice (*Lepeophtheirus salmonis*)^78^. Therefore, shy winners are likely to be at a greater risk for infections and infection-related diseases. Importantly, pyroptosis and *tlr7* may also be linked to the development of autoimmune diseases, despite their primary function in defending against microbial infections^79,80^. While the *viml* gene showed significant overexpression in shy winners compared to bold winners, other genes demonstrated only mild changes in expression, suggesting that the potential for inflammation and pyroptosis should be regarded as a predisposition rather than a definitive outcome for shy winners.

On the contrary, bold winners exhibited signs of iron accumulation in their brain, as indicated by the upregulation of ferritin (*fthl29*), transferrin receptor (*tfr1a*), and the *slc11a2* gene. These genes are implicated in ferroptosis within the KEGG pathways, iron ion transmembrane transporter activity (local network cluster), and abnormal nucleate erythrocyte low saturation (Monarch; STRING database). Transferrin facilitates the endocytosis of iron ions across the blood-brain barrier, allowing for their transport to cells^81^. Excessive free iron ions can lead to oxidative damage in cells due to their role in promoting reactive oxygen species (ROS) production^82^. Ferritin is upregulated in response to increased intracellular iron ion concentrations, serving to store iron in an inactive form^82,83^. The *slc11a2* is associated with iron overload in the brain, although its activity leads to the release and export of stored iron^84^. The upregulation of this gene suggests that bold winners release more iron from storage compared to their shyer counterparts. However, we regard this condition as merely a propensity, as the fold change did not exceed the predefined threshold, with the exception of the *fthl29* gene. However, the upregulation of genes involved in iron metabolism and the consequent iron accumulation has been associated with cognitive impairment, major depressive disorder, type 2 diabetes, and neurodegenerative diseases in humans^73,83,85,86^. Further research in this direction could improve modelling of these diseases in zebrafish models if replication further supports these findings.

*Prodha* gene was found to be upregulated in individuals classified as shy winners, and it has been investigated for its potential variation in relation to different stress coping styles^32^. Its expression increased under conditions of rapid temperature decline in fish^87^ however, it did not exhibit differences between proactive and reactive stress coping styles in another research study^32^. The discrepancy between these findings may be attributed to the observation that only winners displayed variations in the expression of this gene, while losers did not, as evidenced by the current study. Furthermore, this study indicates that although there were significant differences in gene expression, the fold change did not reach the 2-fold threshold. This suggests that additional factors may influence the expression of this gene.

### Repeated social experience

The shy/reactive and bold/proactive coping styles responded differently to social challenge. In bold individuals, the primary difference between losers and winners was observed in the latency to enter the CORN zone, associated with diminished exploratory behaviour in rodents^88^. A similar increase in latency to enter the corner in the MCSF arena was found in rats bred for high alcohol uptake^88^, and in shelter-seeking rats^89^, however, these behavioural changes were accompanied by additional modifications not seen in the current study. Consequently, the social challenge exerted only weak effects on the activity of bold individuals in this investigation. Conversely, substantial alterations in transcriptomics were evident between bold winners and losers. Nine differentially expressed genes, namely *ier2a, ier2b, junba, junbb, fosab, btg2, egr1, nr4a1,* and *npas4a*, showed co-expression in the STRING database and are implicated in significant biological pathways. These differentially expressed genes (DEGs) are associated with the regulation of transcription and DNA-binding activity in the nucleus, as defined by Gene Ontology (GO) classifications for both Molecular Function and Cellular Component (STRING database). The fibroblast growth factor (FGF) signalling pathway involves the *junba, fosab* and *egr1* gene (WikiPathways; STRING) and is regulated by the expression of the *ier2* gene^90^. This signalling pathway is crucial for neuronal growth and differentiation, and it recruits adapter proteins from the ERK1/ERK2 MAPK pathway^91^. The Jun proteins and *fosab* are included among these recruited adapter proteins, functioning downstream of the FGF signalling pathway. Consequently, FGFs may inhibit neuronal cell death and potentially promote oncogenesis if excessively activated^92^, while also facilitating improved spatial learning and neurogenesis^91^. FGFs are essential in the crosstalk between neurons, microglia, and astrocytes^93^. The activity of astrocytes is tightly regulated due to the risk of chronic inflammation, with FGFs helping to maintain their nonreactive state^93^. The upregulation of the *pim1* and *plk1* genes in bold losers in the current study supports the possibility of tumour formation or its predisposition.

Teles et al.^94^ investigated the expression of *fosab* and *egr1* in the social decision-making network (SDMN) within the forebrain and midbrain of fish with varying social experiences. Their findings revealed that the *fosab* gene was significantly upregulated in the preoptic area (POA) of fish identified as losers, compared to those designated as winners. However, no other notable alterations were observed in this study. The *npas4a* gene had been previously shown to be upregulated in the medial part of the dorsal telencephalon of zebrafish even two hours following a winning experience^95^. The current study did not detect these changes in the forebrain or midbrain, potentially due to the dilution of minor effects when analysing broader areas rather than specific nuclei. Nonetheless, when brain samples were collected immediately after the social experience, an upregulation of similar genes that were found in the present study (*npas4a, fosab, nr4a1, ier2*) was observed in losers compared to winners in zebrafish^96^. These genes are believed to relate to the fish’s self-assessment of their social rank,^97^ which aligns with the findings of the present study. Consequently, the bold fish demonstrated a response to their social experience by modifying their brain transcriptome, yet this did not translate into a change in behaviour.

In contrast, shy or reactive fish did not exhibit significant changes in their brain transcriptomes, although their behaviour underwent notable alterations. Specifically, the shy losers manifested an increase in the latency to attack, swam more rapidly through the CIRC zone, and performed higher locomotor activity as evidenced by their movement through the corridors in the zMCSF arena^37^ These behavioural changes in shy losers may indicate elevated anxiety, which can manifest as more intense burst swimming and erratic movements^98,99,100^. Anxiety responses can either involve behavioural disinhibition, such as increased locomotor activity, or inhibition, like freezing^101^. The preference for one type of anxiety response is recognized as a crucial factor in determining stress coping styles^101^. The freezing response is typically the natural reaction to stress in shy or reactive fish^101^ contrasting with the behaviour observed in this study. Additionally, startled fish tend to take longer to approach an opponent and initiate a fight and when they do engage, they employ less aggressive behaviors^102^. The specific types of aggressive behaviours were not assessed in this investigation, but the response of the shy losers seem to resemble the response of startled fish. Conversely, shy winners markedly increased their aggressive actions, indicating a winner effect^18^.

However, only minimal differences in the brain transcriptomes were found between shy winners and losers. This suggests a potential link between the anticipated increased plasticity of shy fish^103^ and their ability to mitigate the impact of social defeat on brain transcriptome changes by altering their behaviour. Behavioural flexibility empowers individuals to modify their actions in response to environmental stimuli ^1^. Thus, the capacity of shy losers to reduce the negative effects of chronic stress from dyadic encounters on their brain transcriptomes may stem from their ability to adjust their behaviour, highlighting a greater level of behavioural flexibility in shy fish within social contexts.

## Limitations of the study

It was not feasible to prevent the occurrence of potential siblings in the current study. The mirror test is a widely utilized non-invasive method for assessing aggression in fish;^104^ however, it is important to recognize that engaging with a mirror image may differ significantly from interacting with a live opponent^105^. Transcriptomic analysis focused on large brain regions (forebrain/midbrain and hindbrain), which limited the ability to conduct detailed examinations of specific brain nuclei. A selection of more defined brain areas may yield a clearer and more comprehensive understanding of our findings.

## Acknowledgements

The behavioural testing was carried out with support of the Uppsala University Behavioural Facility (UUBF), Disciplinary Domain of Medicine and Pharmacy, Uppsala University. The authors gratefully acknowledge the support of the Facias Foundation (P.H. and S.W.), the Swedish Research Council (VR-NT11 2017-03779 to S.W.), and the KI Research Foundation Grants for PhD students (2020-01677, L.B. and P.B.). The authors acknowledge support from the National Genomics Infrastructure in Stockholm funded by Science for Life Laboratory, the Knut and Alice Wallenberg Foundation and the Swedish Research Council, and SNIC/NAISS/Uppsala Multidisciplinary Centre for Advanced Computational Science for assistance with massively parallel sequencing and access to the UPPMAX computational infrastructure. We also would like to thank the core facility at NEO, BEA, Bioinformatics and Expression Analysis, which is supported by the board of research at the Karolinska Institute and the research committee at the Karolinska hospital. We would like to thank to Per-Ove Thörnqvist, Erika Roman, and Laura Vossen for their consultations throughout this experiment.

## Author contributions

Conceptualization, S.W. and P.B.; Methodology, J.A, O.S.S, and S.W; Formal Analysis, P.H. and D.B.; Investigation, P.H., L.B., and D.B; Resources, S.W. and P.B., Data Curation, D.B.; Writing – Original Draft, P.H.; Writing – Review & Editing, L.B., D.B., J.A., O.S.S., S.W., and P.B.; Visualization, L.B. and D.B.; Supervision, S.W. and P.B.; Funding Acquisition, S.W. and P.B.

## Data and code availability

The data reported in this paper will be shared by the lead contact upon request. Any additional information required to reanalyse the data reported in this paper is available from the lead contact upon request.

## Declaration of generative AI and AI-assisted technologies

During the preparation of this work the author(s) used Avidnote in order to improve readability of the text. After using this tool/service, the author(s) reviewed and edited the content as needed and take(s) full responsibility for the content of the published article.

## Supplemental information titles and legends

SF-1 – Artificial Selection Program

SF-2 – Behavioural analysis

SF-3 – Differentially expressed genes from the transcriptomic analysis

## References

1. Coppens CM, De Boer SF, Koolhaas JM. Coping styles and behavioural flexibility: Towards underlying mechanisms. Philosophical Transactions of the Royal Society B: Biological Sciences 2010, 365:4021–4028.

2. Sørensen C, Johansen IB, Øverli Ø. Neural plasticity and stress coping in teleost fishes. Gen Comp Endocrinol 2013, 181:25–34.

3. Oliveira RF. Social plasticity in fish: Integrating mechanisms and function. J Fish Biol 2012, 81:2127–2150.

4. Benus RF, Bohus B, Koolhaas JM, van Oortmerssen GA. Heritable variation for aggression as a reflection of individual coping strategies. Experientia 1991, 47:1008–1019.

5. de Boer SF, Buwalda B, Koolhaas JM. Untangling the neurobiology of coping styles in rodents: Towards neural mechanisms underlying individual differences in disease susceptibility. Neurosci Biobehav Rev 2017, 74:401–422.

6. Ruiz-Gomez M de L, Huntingford FA, Øverli Ø, Thörnqvist PO, Höglund E. Response to environmental change in rainbow trout selected for divergent stress coping styles. Physiol Behav 2011, 102:317–322.

7. Winberg S, Thörnqvist PO. Role of brain serotonin in modulating fish behaviour. Curr Zool 2016, 62:317–323.

8. Zupanc GKH, Lamprecht J. Towards a Cellular Understanding of Motivation: Structural Reorganization and Biochemical Switching as Key Mechanisms of Behavioral Plasticity. Ethology 2000, 106:467–477.

9. Dhuriya YK, Sharma D. Neuronal Plasticity: Neuronal Organization is Associated with Neurological Disorders. Journal of Molecular Neuroscience 2020, 70:1684–1701.

10. Koolhaas JM, de Boer SF, Coppens CM, Buwalda B. Neuroendocrinology of coping styles: Towards understanding the biology of individual variation. Front Neuroendocrinol 2010, 31:307–321.

11. Castanheira MF, Conceição LEC, Millot S, et al. Coping styles in farmed fish: consequences for aquaculture. Rev Aquac 2017, 9:23–41.

12. Vindas MA, Gorissen M, Höglund E, et al. How do individuals cope with stress? Behavioural, physiological and neuronal differences between proactive and reactive coping styles in fish. Journal of Experimental Biology 2017, 220:1524–1532.

13. Laursen DC, L. Olsén H, Ruiz-Gomez M de L, Winberg S, Höglund E. Behavioural responses to hypoxia provide a non-invasive method for distinguishing between stress coping styles in fish. Appl Anim Behav Sci 2011, 132:211–216.

14. Puglisi-Allegra S, Andolina D. Serotonin and stress coping. Behavioural Brain Research 2015, 277:58–67.

15. Ebner K, Singewald GM, Whittle N, Ferraguti F, Singewald N. Neurokinin 1 receptor antagonism promotes active stress coping via enhanced septal 5-HT transmission. Neuropsychopharmacology 2008, 33:1929–1941.

16. Frost AJ, Winrow-Giffen A, Ashley PJ, Sneddon LU. Plasticity in animal personality traits: Does prior experience alter the degree of boldness? Proceedings of the Royal Society B: Biological Sciences 2007, 274:333–339.

17. Timmer M, Sandi C. A role for glucocorticoids in the long-term establishment of a social hierarchy. Psychoneuroendocrinology 2010, 35:1543–1552.

18. Oliveira RF, Silva JF, Simões JM. Fighting zebrafish: Characterization of aggressive behavior and winner-loser effects. Zebrafish 2011, 8:73–81.

19. Li CY, Tseng YC, Chen YJ, Yang Y, Hsu Y. Personality and physiological traits predict contest interactions in *Kryptolebias marmoratus*. Behavioural Processes 2020, 173:104079.

20. Perrone R, Silva A. Vasotocin increases dominance in the weakly electric fish *Brachyhypopomus gauderio*. J Physiol 2016, 110:119–126.

21. Weger M, Sevelinges Y, Grosse J, de Suduiraut IG, Zanoletti O, Sandi C. Increased brain glucocorticoid actions following social defeat in rats facilitates the long-term establishment of social subordination. Physiol Behav 2018, 186:31–36.

22. Coppens CM, Sipornmongcolchai T, Wibrand K, et al. Social defeat during adolescence and adulthood differentially induce *BDNF*-regulated immediate early genes. Front Behav Neurosci 2011, 5:1–8.

23. Wallace KJ, Choudhary KD, Kutty LA, et al. Social ascent changes cognition, behaviour and physiology in a highly social cichlid fish. Philosophical Transactions of the Royal Society B: Biological Sciences 2022 377:20200448.

24. Oliveira RF. Social behavior in context: Hormonal modulation of behavioural plasticity and social competence. Integr Comp Biol 2009, 49:423–440.

25. Pavlidis M, Sundvik M, Chen YC, Panula P. Adaptive changes in zebrafish brain in dominant- subordinate behavioural context. Behavioural Brain Research 2011, 225:529–537.

26. Camerlink I, Arnott G, Farish M, Turner SP. Complex contests and the influence of aggressiveness in pigs. Anim Behav 2016, 121:71–78.

27. Pravosudov V V., Mendoza SP, Clayton NS. The relationship between dominance, corticosterone, memory, and food caching in mountain chickadees (*Poecile gambeli*). Horm Behav 2003, 44:93–102.

28. Spritzer MD, Meikle DB, Solomon NG. The relationship between dominance rank and spatial ability among male meadow voles (*Microtus pennsylvanicus)*. J Comp Psychol 2004, 118:332–339.

29. Barnard CJ, Luo N. Acquisition of dominance status affects maze learning in mice. Behavioural Processes 2002, 60:53–59.

30. Sneddon LU, Hawkesworth S, Braithwaite VA, Yerbury J. Impact of environmental disturbance on the stability and benefits of individual status within dominance hierarchies. Ethology 2006, 112:437–447.

31. Frost AJ, Thomson JS, Smith C, et al. Environmental change alters personality in the rainbow trout, *Oncorhynchus mykiss*. Anim Behav 2013, 85:1199–1207.

32. Wong RY, Lamm MS, Godwin J. Characterizing the neurotranscriptomic states in alternative stress coping styles. BMC Genomics 2015, 16:1–11.

33. Alfonso S, Zupa W, Manfrin A, et al. Stress coping styles: Is the basal level of stress physiological indicators linked to behaviour of sea bream? Appl Anim Behav Sci 2020, 231:105085.

34. Alfonso S, Sadoul B, Gesto M, et al. Coping styles in European sea bass: The link between boldness, stress response and neurogenesis. Physiol Behav 2019, 207:76–85.

35. Mu MD, Geng HY, Rong KL, et al. A limbic circuitry involved in emotional stress-induced grooming. Nat Commun 2020, 11:1–16.

36. Kalueff A V., Tuohimaa P. Contrasting grooming phenotypes in three mouse strains markedly different in anxiety and activity (129S1, BALB/c and NMRI). Behavioural Brain Research 2005, 160:1–10.

37. Vossen LE, Brunberg R, Rådén P, Winberg S, Roman E. The zebrafish Multivariate Concentric Square Field: A Standardized Test for Behavioral Profiling of Zebrafish (*Danio rerio*). Front Behav Neurosci 2022, 16:744533.

38. Friard O, Gamba M. BORIS: a free, versatile open-source event-logging software for video/audio coding and live observations. Methods Ecol Evol 2016, 7:1325–1330.

39. R Core Team. R: A language and environment for statistical computing. Published online 2022.

40. Peterson RA. Finding Optimal Normalizing Transformations via bestNormalize. R J 2021, 13:310–329.

41. Wickham H, Chang W. Package ggplot2: An Implementation of the Grammar of Graphics. *CRAN Repository* 2014, 1-189.

42. Russel L. emmeans: estimated Marginal Means, aka Least-Squares Means. *R package version 1.2* 2019.

43. Bates D, Mächler M, Bolker BM, Walker SC. Fitting linear mixed-effects models using lme4. J Stat Softw 2015, 67:1–51.

44. Fox J, Weisberg S, Bates D, et al. The car Package. R Foundation for Statistical computing 2012, 1109:1431.

45. Ripely B, Venables B, Douglas BM, Hornik K, Gebhardt A, Firth D. Package ‘MASS’: Functions and datasets to support Venables and Ripley. Modern Applied Statistics with S. CRAN Repository 2023, 1-164.

46. Huntingford FA, Andrew G, Mackenzie S, et al. Coping strategies in a strongly schooling fish, the common carp *Cyprinus carpio*. J Fish Biol 2010, 76:1576–1591.

47. Jenjan H, Mesquita F, Huntingford F, Adams C. Respiratory function in common carp with different stress coping styles: A hidden cost of personality traits? Anim Behav 2013, 85:1245–1249.

48. Tudorache C, Ter Braake A, Tromp M, Slabbekoorn H, Schaaf MJM. Behavioural and physiological indicators of stress coping styles in larval zebrafish. Stress 2015, 18:121–128.

49. Linares-Cordova JF, Rey-Planellas S, Boglino A, et al. Flathead grey mullet (*Mugil cephalus*) juveniles exhibit consistent proactive and reactive stress coping styles. Aquaculture 2024, 578:740012.

50. Øverli Ø, Sørensen C, Pulman KGT, et al. Evolutionary background for stress-coping styles: Relationships between physiological, behavioral, and cognitive traits in non-mammalian vertebrates. Neurosci Biobehav Rev 2007, 31:396–412.

51. Laursen DC, L. Olsén H, Ruiz-Gomez M de L, Winberg S, Höglund E. Behavioural responses to hypoxia provide a non-invasive method for distinguishing between stress coping styles in fish. Appl Anim Behav Sci 2011, 132:211–216.

52. Bergstedt JH, Pfalzgraff T, Skov P V. Hypoxia tolerance and metabolic coping strategies in *Oreochromis niloticus*. Comp Biochem Physiol A Mol Integr Physiol 2021, 257:110956.

53. Castanheira MF, Conceição LEC, Millot S, et al. Coping styles in farmed fish: consequences for aquaculture. Rev Aquac 2017, 9:23–41.

54. Jeffrey JD, Esbaugh AJ, Vijayan MM, Gilmour KM. Modulation of hypothalamic-pituitary- interrenal axis function by social status in rainbow trout. Gen Comp Endocrinol 2012, 176:201–210.

55. Tort L. Stress and immune modulation in fish. Dev Comp Immunol 2011, 35:1366–1375.

56. Winberg S, Sneddon L. Impact of intraspecific variation in teleost fishes: aggression, dominance status and stress physiology. Journal of Experimental Biology 2022, 225:jeb169250.

57. Culbert BM, Gilmour KM. Rapid recovery of the cortisol response following social subordination in rainbow trout. Physiol Behav 2016, 164:306–313.

58. Sukhareva E V. The role of the corticotropin-releasing hormone and its receptors in the regulation of stress response. Vavilovskii Zhurnal Genet Selektsii 2021, 25:216–223.

59. Kyjacova L, Saup R, Rönsch K, et al. *IER2*-induced senescence drives melanoma invasion through osteopontin. Oncogene 2021, 40:6494–6512.

60. Rossi JJ, Rosenfeld JA, Chan KM, et al. Molecular characterisation of rare loss-of-function *NPAS3* and *NPAS4* variants identified in individuals with neurodevelopmental disorders. Sci Rep 2021, 11:1–11.

61. Deng S, Chen B, Huo J, Liu X. Therapeutic potential of *NR4A1* in cancer: Focus on metabolism. Front Oncol 2022, 12:1–13.

62. Hughes BW, Siemsen BM, Tsvetkov E, et al. *NPAS4* in the medial prefrontal cortex mediates chronic social defeat stress-induced anhedonia-like behaviour and reductions in excitatory synapses. Elife 2023, 12:1–28.

63. Lissek T, Andrianarivelo A, Saint-Jour E, et al. *Npas4* regulates medium spiny neuron physiology and gates cocaine-induced hyperlocomotion. EMBO Rep 2021, 22.

64. Brivio P, Gallo MT, Gruca P, et al. Chronic N-Acetyl-Cysteine Treatment Enhances the Expression of the Immediate Early Gene *Nr4a1* in Response to an Acute Challenge in Male Rats: Comparison with the Antidepressant Venlafaxine. Int J Mol Sci 2023, 24.

65. Øverli Ø, Korzan WJ, Höglund E, et al. Stress coping style predicts aggression and social dominance in rainbow trout. Horm Behav 2004, 45:235–241.

66. Ji C, Zhao H, Li D, et al. Role of *Wdr45b* in maintaining neural autophagy and cognitive function. Autophagy 2020, 16:615–625.

67. Yang L, Zeng W, Sun H, et al. Bioinformatical Analysis of Gene Expression Omnibus Database Associates *TAF7/CCNB1, TAF7/CCNA2*, and *GTF2E2/CDC20* Pathways with Glioblastoma Development and Prognosis. World Neurosurg 2020, 138:492-514.

68. Seifert C, Balz E, Herzog S, et al. *Pim1* inhibition affects glioblastoma stem cell behaviour and kills glioblastoma stem-like cells. Int J Mol Sci 2021, 22:11126.

69. Ye M, Wang S, Qie JB, Sun PL. SPRY4-AS1, A Novel Enhancer RNA, Is a Potential Novel Prognostic Biomarker and Therapeutic Target for Hepatocellular Carcinoma. Front Oncol 2021, 11:1–9.

70. Peng Y, Liu Y, Zheng R, et al. *PLK1* maintains DNA methylation and cell viability by regulating phosphorylation-dependent *UHRF1* protein stability. Cell Death Discov 2023, 9:1–13.

71. Acevedo A, Torres F, Kiwi M, et al. Metabolic switch in the aging astrocyte supported via integrative approach comprising network and transcriptome analyses. Aging 2023, 19:9896.

72. Paulin D, Lilienbaum A, Kardjian S, Agbulut O, Li Z. Vimentin: Regulation and pathogenesis. Biochimie 2022, 197:96–112.

73. Chen KZ, Liu SX, Li YW, et al. Vimentin as a potential target for diverse nervous system diseases. Neural Regen Res 2023, 18:969–975.

74. Xun Y, Yang H, Kaminska B, You H. Toll-like receptors and toll-like receptor-targeted immunotherapy against glioma. J Hematol Oncol 2021, 14:1–32.

75. Yu P, Zhang X, Liu N, Tang L, Peng C, Chen X. Pyroptosis: mechanisms and diseases. Signal Transduct Target Ther 2021, 6:128.

76. Wen H, Guo D, Zhao Z, et al. Novel pyroptosis-associated genes signature for predicting the prognosis of sarcoma and validation. Biosci Rep 2022, 42:1–13.

77. Pickering AD, Pottinger TG. Stress responses and disease resistance in salmonid fish: Effects of chronic elevation of plasma cortisol. Fish Physiol Biochem 1989, 7:253–258.

78. Kittilsen S, Johansen IB, Braastad BO, Øverli Ø. Pigments, parasites and personality: Towards a unifying role for steroid hormones? PLoS One 2012, 7:26–29.

79. David C, Badonyi M, Kechiche R, et al. Interface Gain-of-Function Mutations in *TLR7* Cause Systemic and Neuro-inflammatory Disease. J Clin Immunol 2024, 44:1–9.

80. Burdette BE, Esparza AN, Zhu H, Wang S. Gasdermin D in pyroptosis. Acta Pharm Sin B 2021, 11:2768–2782.

81. Thomsen MS, Johnsen KB, Kucharz K, Lauritzen M, Moos T. Blood–Brain Barrier Transport of Transferrin Receptor-Targeted Nanoparticles. Pharmaceutics 2022, 14:1–17.

82. Zhang N, Yu X, Xie J, Xu H. New Insights into the Role of Ferritin in Iron Homeostasis and Neurodegenerative Diseases. Mol Neurobiol 2021, 58:2812–2823.

83. Urrutia PJ, Bórquez DA, Núñez MT. Inflaming the brain with iron. Antioxidants 2021, 10:1-27.

84. Volk Robertson K, Schleh MW, Harrison FE, Hasty AH. Microglial-specific knockdown of iron import gene, *Slc11a2*, blunts LPS-induced neuroinflammatory responses in a sex-specific manner. Brain Behav Immun 2024, 116:370-384.

85. Hoshi K, Ito H, Abe E, et al. Transferrin biosynthesized in the brain is a novel biomarker for Alzheimer’s disease. Metabolites 2021, 11:616.

86. Chang X, Ma M, Chen L, et al. Identification and Characterization of Elevated Expression of Transferrin and Its Receptor *TfR1* in Mouse Models of Depression. Brain Sci 2022, 12:1–17.

87. Liu Z, Zhu L, Wang X, et al. Application of transcriptome analysis to investigate the effects of long-term low temperature stress on liver function in the tiger puffer (*Takifugu rubripes*). Front Mar Sci 2022, 9:1-15.

88. Roman E, Stewart RB, Bertholomey ML, et al. Behavioural profiling of multiple pairs of rats selectively bred for high and low alcohol intake using the MCSF test. Addiction Biology 2012, 17:33–46.

89. Lundberg S, Högman C, Roman E. Adolescent exploratory strategies and behavioural types in the multivariate concentric square field TM test. Front Behav Neurosci 2019, 13:1–18.

90. Hong SK, Dawid IB. FGF-dependent left-right asymmetry patterning in zebrafish is mediated by *Ier2* and *Fibp1*. Proc Natl Acad Sci USA 2009, 106:2230–2235.

91. Klimaschewski L, Claus P. Fibroblast Growth Factor Signalling in the Diseased Nervous System. Mol Neurobiol 2021, 58:3884–3902.

92. Ardizzone A, Scuderi SA, Giuffrida D, et al. Role of fibroblast growth factors receptors (*FGFR*s) in brain tumours, focus on astrocytoma and glioblastoma. Cancers 2020, 12:1–22.

93. Kang W, Balordi F, Su N, Chen L, Fishell G, Hébert JM. Astrocyte activation is suppressed in both normal and injured brain by *FGF* signalling. Proc Natl Acad Sci USA 2014, 111:2987–2995.

94. Teles MC, Almeida O, Oliveira RF. Social interactions elicit rapid shifts in functional connectivity in the social decision-making network of zebrafish. Proc Roy Soc B: Biol Sci 2015, 282:20151099.

95. Teles MC, Cardoso SD, Oliveira RF. Social plasticity relies on different neuroplasticity mechanisms across the brain social decision-making network in zebrafish. Front Behav Neurosci 2016, 10:16.

96. Oliveira RF, Simes JM, Teles MC, Oliveira CR, Becker JD, Lopes JS. Assessment of fight outcome is needed to activate socially driven transcriptional changes in the zebrafish brain. Proc Natl Acad Sci USA 2016, 113:654–661.

97. Oliveira RF, Simes JM, Teles MC, Oliveira CR, Becker JD, Lopes JS. Assessment of fight outcome is needed to activate socially driven transcriptional changes in the zebrafish brain. Proc Natl Acad Sci USA 2016, 113:654–661.

98. Maximino C, de Brito TM, da Silva Batista AW, Herculano AM, Morato S, Gouveia A. Measuring anxiety in zebrafish: A critical review. Behavioural Brain Research 2010, 214:157–171.

99. Caldarone BJ, King SL, Picciotto MR. Sex differences in anxiety-like behavior and locomotor activity following chronic nicotine exposure in mice. Neurosci Lett 2008, 439:187–191.

100. Tran S, Nowicki M, Muraleetharan A, Chatterjee D, Gerlai R. Neurochemical factors underlying individual differences in locomotor activity and anxiety-like behavioral responses in zebrafish. Prog Neuropsychopharmacol Biol Psychiatry 2016, 65:25–33.

101. Koolhaas JM, Korte SM, De Boer SF, et al. Coping styles in animals: Current status in behavior and stress- physiology. Neurosci Biobehav Rev 1999, 23:925–935.

102. Arnott G, Elwood R. Probing aggressive motivation in a cichlid fish. Biol Lett 2009, 5:762–764.

103. Ruiz-Gomez MDL, Kittilsen S, Höglund E, et al. Behavioral plasticity in rainbow trout (*Oncorhynchus mykiss*) with divergent coping styles: When doves become hawks. Horm Beh 2008, 54:534–538.

104. Way GP, Ruhl N, Snekser JL, Kiesel AL, McRobert SP. A comparison of methodologies to test aggression in zebrafish. Zebrafish 2015, 12:144–151.

105. Balzarini V, Taborsky M, Wanner S, Koch F, Frommen JG. Mirror, mirror on the wall: The predictive value of mirror tests for measuring aggression in fish. Behav Ecol Sociobiol 2014, 68:871–878.

