## Supplementary material for "Effects of stress coping styles and social defeat on zebrafish behaviour and brain transcriptomics": SF-1

*Artificial selection*

Selective breeding for boldness was used to generate lines of zebrafish manifesting divergent behavioural phenotype. The founder population were wild-caught zebrafish captured in West Bengal, India. Their offspring (n = 1000) spawned on site were transported to Norwegian Technical University (Trondheim, Norway). Eggs collected from the offspring of wild-caught zebrafish, the $F_0$ generation, were transported to Uppsala University (Uppsala, Sweden; curtesy of Dr. Fredrik Jutfelt). The adult fish ($F_0$) were tagged by p-Chip (0.5 × 0.5 × 0.1 mm) according to p-Chip Implantation Protocol (Chen et al., 2017), except for the fact that 200 mg/L of benzocaine (Merck KGaA, Darmstadt, Germany) were used for anaesthesia in the present study. The fish were consequently tested twice in the novel tank diving (NTD) test for boldness (latency to enter the top zone of the tank) according to Thörnqvist et al. (2019). The fish (250-600 individuals) were selected from the upper and lower extremes of the population, i.e., the fish being boldest and the fish being the shyest. Proactive (bold) and reactive (shy) lines were generated in duplicate (i.e., R1 shy, R2 shy, R1 bold, R2 bold) and randomly mated within the same selected line. Parents (125-300 fish) were selected for mating of the $F_1$ generation. The same process was repeated to generate the $F_2$ generation. The $F_2$ generation of fish significantly differed in boldness in the NTD test (see **SF-IA**).

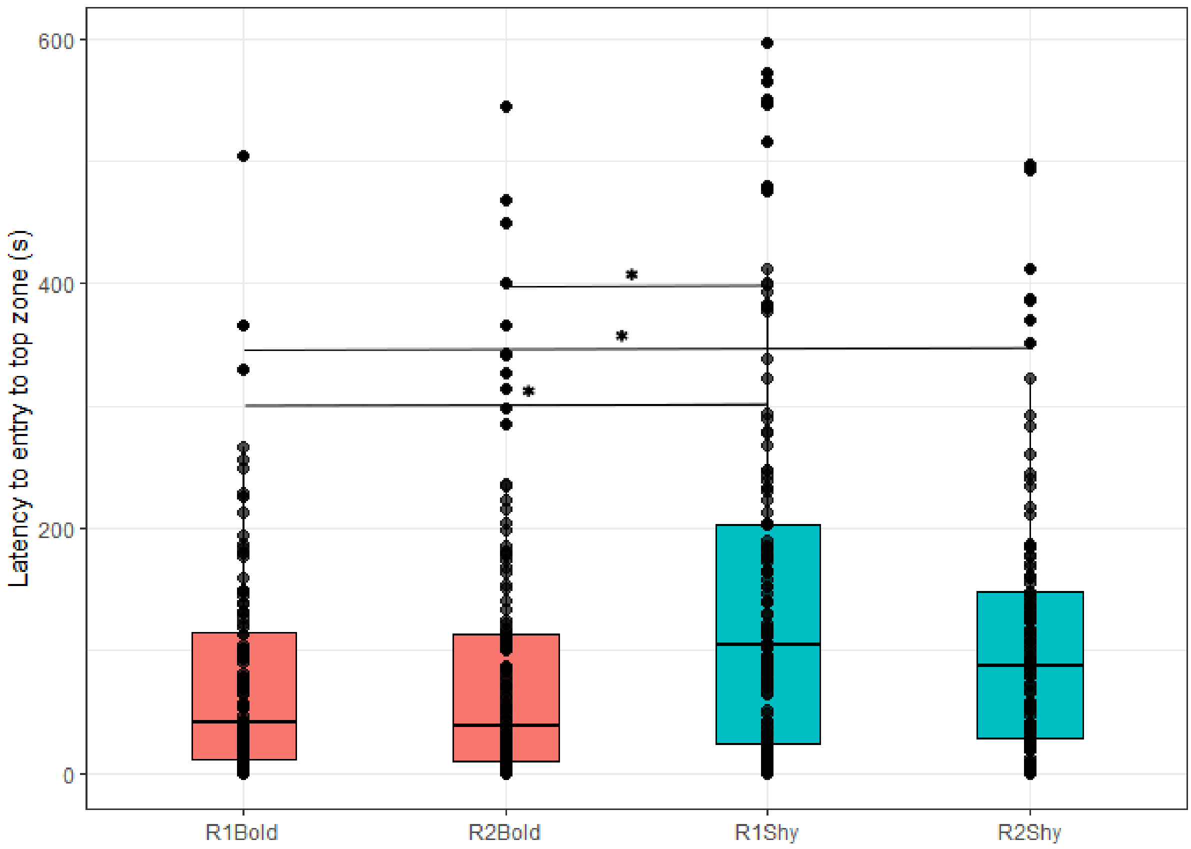


**SF-1A**: The differences in the latency to enter the top zone between bold and shy selected lines generated in duplicate (R1, R2 lines for both shy and bold fish) of the $F_2$ generation of fish. Ordernorm transformation was used to normalize the data. Linear mixed effect models with Line as fixed effect and individual as a random effect were applied to detect significant differences at $\alpha = 0.05$ and are highlighted as asterisk. Emmeans function was as a pairwise comparison between selected lines.

**References**

Chen, C.-H., Poss, K., & Hughes, H. (2017). Adult Zebrafish p-Chip Implantation Protocol. *PharmaSeq*.

Thörnqvist, P. O., McCarrick, S., Ericsson, M., Roman, E., & Winberg, S. (2019). Bold zebrafish (*Danio rerio*) express higher levels of delta opioid and dopamine D2 receptors in the brain compared to shy fish. *Behavioural Brain Research*, 359: 927–934.
