## Supplementary material for "Effects of stress coping styles and social defeat on zebrafish behaviour and brain transcriptomics": SF-2

**SF-2**: Results of the contrast analysis using the "emmeans" package in R from the Mirror Test (MT) and the zebrafish Multivariate Concentric Square Field (zMCSF) test. The non-normal data were transformed based on the recommendation of the "bestNormalize" package in R. 1 = the first trial of MT, 1-2 = the percentage change from the first to the second trial of MT, B = bold line, BL = bold loser, BW = bold winner, NB = negative binomial distribution, oN = orderNorm transformation, PD = Poisson distribution; S = shy line, SL = shy loser, sqrt = square-root transformation, SW = shy winner, YJ = Yeo Johnson transformation.

| **Comparisons** | **Test** | **df** | **Test statistics value** | $p_{adj}$ |
|---|---|---|---|---|
| **MT – Number of aggressive interactions** | | | | |
| B (1) – S (1) | GLMM (PD) | | 2.78 | .005** |
| BL (1-2) – BW (1-2) | ANOVA (oN) | 28 | 1.44 | .160 |
| SL (1-2) – SW (1-2) | ANOVA (oN) | 28 | -1.03 | .0312 |
| BL (1-2) – SL (1-2) | ANOVA (oN) | 28 | -0.16 | .875 |
| BW (1-2) – SW (1-2) | ANOVA (oN) | 28 | -2.63 | .014* |
| **MT – Latency to the first attack** | | | | |
| B (1) – S (1) | ANOVA (log) | 30 | 0.65 | .520 |
| BL (1-2) – BW (1-2) | ANOVA (oN) | 28 | -0.92 | .365 |
| SL (1-2) – SW (1-2) | ANOVA (oN) | 28 | 1.38 | .180 |
| BL (1-2) – SL (1-2) | ANOVA (oN) | 28 | -2.28 | .031* |
| BW (1-2) – SW (1-2) | ANOVA (oN) | 28 | 0.02 | .985 |
| **MT – Total duration of attack** | | | | |
| B (1) – S (1) | ANOVA (oN) | 30 | 2.71 | .011* |
| BL (1-2) – BW (1-2) | ANOVA (oN) | 28 | 0.08 | .934 |
| SL (1-2) – SW (1-2) | ANOVA (oN) | 28 | -0.64 | .529 |
| BL (1-2) – SL (1-2) | ANOVA (oN) | 28 | 0.18 | .860 |
| BW (1-2) – SW (1-2) | ANOVA (oN) | 28 | -0.54 | .592 |
| **MT – Average duration of attack** | | | | |
| B (1) – S (1) | ANOVA (oN) | 30 | 2.45 | .020* |
| BL (1-2) – BW (1-2) | ANOVA (oN) | 28 | -0.84 | .411 |
| SL (1-2) – SW (1-2) | ANOVA (oN) | 28 | 0.04 | .970 |
| BL (1-2) – SL (1-2) | ANOVA (oN) | 28 | -0.50 | .624 |
| BW (1-2) – SW (1-2) | ANOVA (oN) | 28 | 0.38 | .709 |
| **MT – Number of displaced behaviours** | | | | |
| B (1) – S (1) | GLMM (NB) | | -0.40 | .690 |
| BL (1-2) – BW (1-2) | ANOVA (oN) | 28 | 0.55 | .584 |
| SL (1-2) – SW (1-2) | ANOVA (oN) | 28 | 1.56 | .131 |
| BL (1-2) – SL (1-2) | ANOVA (oN) | 28 | -0.33 | .745 |
| BW (1-2) – SW (1-2) | ANOVA (oN) | 28 | 0.67 | .507 |
| **MT – Latency to the first displaced behaviour** | | | | |
| B (1) – S (1) | ANOVA (oN) | 30 | -0.87 | .391 |
| BL (1-2) – BW (1-2) | ANOVA (oN) | 28 | 0.04 | .981 |
| SL (1-2) – SW (1-2) | ANOVA (oN) | 28 | -0.48 | .637 |
| BL (1-2) – SL (1-2) | ANOVA (oN) | 28 | 0.42 | .676 |
| BW (1-2) – SW (1-2) | ANOVA (oN) | 28 | -0.08 | .938 |
| **MT – Total duration of displaced behaviour** | | | | |
| B (1) – S (1) | ANOVA (oN) | 30 | -0.49 | .631 |
| BL (1-2) – BW (1-2) | ANOVA (oN) | 28 | -0.19 | .853 |
| SL (1-2) – SW (1-2) | ANOVA (oN) | 28 | 1.15 | .260 |
| BL (1-2) – SL (1-2) | ANOVA (oN) | 28 | -0.14 | .892 |
| BW (1-2) – SW (1-2) | ANOVA (oN) | 28 | 1.20 | .241 |
| **MT – Average duration of displaced behaviour** | | | | |
| B (1) – S (1) | ANOVA (oN) | 30 | -0.65 | .523 |
| BL (1-2) – BW (1-2) | ANOVA (oN) | 28 | -0.69 | .496 |
| SL (1-2) – SW (1-2) | ANOVA (oN) | 28 | 0.52 | .610 |
| BL (1-2) – SL (1-2) | ANOVA (oN) | 28 | 0.33 | .747 |
| BW (1-2) – SW (1-2) | ANOVA (oN) | 28 | 1.53 | .137 |

| | | | | |
|---|---|---|---|---|
| **zMCSF – Total distance moved (arena)** | | | | |
| BL - BW | ANOVA | 28 | -0.18 | .861 |
| BL - SL | ANOVA | 28 | 1.16 | .256 |
| BW - SW | ANOVA | 28 | 0.99 | .333 |
| SL - SW | ANOVA | 28 | -0.35 | .728 |
| **zMCSF – Average velocity (arena)** | | | | |
| BL - BW | ANOVA | 28 | -0.25 | .808 |
| BL - SL | ANOVA | 28 | 0.99 | .332 |
| BW - SW | ANOVA | 28 | 1.01 | .323 |
| SL - SW | ANOVA | 28 | -0.23 | .822 |
| **zMCSF – Total activity (arena)** | | | | |
| BL - BW | ANOVA | 28 | -0.28 | .783 |
| BL - SL | ANOVA | 28 | 1.13 | .269 |
| BW - SW | ANOVA | 28 | 1.15 | .260 |
| SL - SW | ANOVA | 28 | -0.26 | .800 |
| **zMCSF – Immobility (arena)** | | | | |
| BL - BW | ANOVA (sqrt) | 28 | -0.37 | .715 |
| BL - SL | ANOVA (sqrt) | 28 | -1.00 | .326 |
| BW - SW | ANOVA (sqrt) | 28 | 0.48 | .638 |
| SL - SW | ANOVA (sqrt) | 28 | 1.11 | .278 |
| **zMCSF – Total distance moved (zones)** | | | | |
| START: BL – BW | LMM (YJ) | 180 | 0.32 | .751 |
| START: BL - SL | LMM (YJ) | 180 | 0.84 | .401 |
| START: BW - SW | LMM (YJ) | 171 | 0.29 | .773 |
| START: SL – SW | LMM (YJ) | 171 | -0.25 | .802 |
| DCR: BL – BW | LMM (YJ) | 180 | 0.21 | .832 |
| DCR: BL - SL | LMM (YJ) | 180 | -0.31 | .759 |
| DCR: BW - SW | LMM (YJ) | 171 | -0.84 | .403 |
| DCR: SL – SW | LMM (YJ) | 171 | -0.30 | .762 |
| CORR1: BL – BW | LMM (YJ) | 171 | -0.66 | .507 |
| CORR1: BL - SL | LMM (YJ) | 195 | 0.14 | .888 |
| CORR1: BW - SW | LMM (YJ) | 171 | 1.56 | .120 |
| CORR1: SL – SW | LMM (YJ) | 195 | 0.70 | .484 |
| CORN: BL – BW | LMM (YJ) | 180 | -0.92 | .359 |
| CORN: BL - SL | LMM (YJ) | 218 | -0.20 | .845 |
| CORN: BW - SW | LMM (YJ) | 171 | 2.15 | .033* |
| CORN: SL – SW | LMM (YJ) | 211 | 1.28 | .203 |
| CORR2: BL – BW | LMM (YJ) | 180 | -1.02 | .312 |
| CORR2: BL - SL | LMM (YJ) | 202 | 0.01 | .993 |
| CORR2: BW - SW | LMM (YJ) | 171 | 2.53 | .012* |
| CORR2: SL – SW | LMM (YJ) | 195 | 1.39 | .166 |
| RAMP1: BL – BW | LMM (YJ) | 180 | 0.31 | .759 |
| RAMP1: BL - SL | LMM (YJ) | 202 | 0.71 | .478 |
| RAMP1: BW - SW | LMM (YJ) | 171 | 1.56 | .122 |
| RAMP1: SL - SW | LMM (YJ) | 195 | 1.03 | .304 |
| RAMP2: BL – BW | LMM (YJ) | 180 | 0.53 | .596 |
| RAMP2: BL - SL | LMM (YJ) | 190 | 2.45 | .015* |
| RAMP2: BW - SW | LMM (YJ) | 171 | 0.20 | .841 |
| RAMP2: SL – SW | LMM (YJ) | 182 | -1.79 | .075 |
| RAMP3: BL – BW | LMM (YJ) | 180 | 0.13 | .898 |
| RAMP3: BL - SL | LMM (YJ) | 180 | 2.58 | .011* |
| RAMP3: BW - SW | LMM (YJ) | 171 | 0.89 | .374 |
| RAMP3: SL – SW | LMM (YJ) | 171 | -1.63 | .105 |
| RAMP4: BL – BW | LMM (YJ) | 190 | -1.58 | .116 |
| RAMP4: BL - SL | LMM (YJ) | 180 | 1.68 | .094 |
| RAMP4: BW - SW | LMM (YJ) | 182 | 3.59 | < .001*** |
| RAMP4: SL - SW | LMM (YJ) | 171 | 0.29 | .771 |
| CIRC: BL – BW | LMM (YJ) | 203 | 0.20 | .845 |
| CIRC: BL – SL | LMM (YJ) | 190 | 0.55 | .586 |
| CIRC: BW – SW | LMM (YJ) | 204 | 0.01 | .996 |

| | | | | |
|---|---|---|---|---|
| CIRC: SL – SW | LMM (YJ) | 192 | -0.34 | .735 |
| CENT: BL – BW | LMM (YJ) | 171 | 0.62 | .539 |
| CENT: BL – SL | LMM (YJ) | 171 | 1.57 | .119 |
| CENT: BW – SW | LMM (YJ) | 171 | -0.49 | .623 |
| CENT: SL – SW | LMM (YJ) | 171 | -1.44 | .151 |
| REST: BL – BW | LMM (YJ) | 171 | -0.03 | .976 |
| REST: BL – SL | LMM (YJ) | 171 | 0.07 | .946 |
| REST: BW - SW | LMM (YJ) | 171 | 0.18 | .856 |
| REST: SL - SW | LMM (YJ) | 171 | 0.09 | .932 |
| **zMCSF – Average velocity (zones)** | | | | |
| START: BL – BW | LMM (oN) | 119 | -0.88 | .382 |
| START: BL - SL | LMM (oN) | 119 | 1.18 | .239 |
| START: BW - SW | LMM (oN) | 111 | 1.54 | .126 |
| START: SL – SW | LMM (oN) | 111 | -0.57 | .570 |
| DCR: BL – BW | LMM (oN) | 119 | -0.07 | .947 |
| DCR: BL - SL | LMM (oN) | 119 | 0.69 | .491 |
| DCR: BW - SW | LMM (oN) | 111 | 0.44 | .659 |
| DCR: SL – SW | LMM (oN) | 111 | -0.34 | .738 |
| CORR1: BL – BW | LMM (oN) | 111 | 0.26 | .795 |
| CORR1: BL - SL | LMM (oN) | 129 | 0.57 | .568 |
| CORR1: BW - SW | LMM (oN) | 111 | -0.14 | .888 |
| CORR1: SL – SW | LMM (oN) | 129 | -0.46 | .647 |
| CORN: BL – BW | LMM (oN) | 119 | -0.15 | .880 |
| CORN: BL - SL | LMM (oN) | 150 | -0.50 | .621 |
| CORN: BW - SW | LMM (oN) | 111 | -1.21 | .230 |
| CORN: SL – SW | LMM (oN) | 143 | -0.74 | .459 |
| CORR2: BL – BW | LMM (oN) | 119 | -0.27 | .784 |
| CORR2: BL - SL | LMM (oN) | 136 | 0.21 | .837 |
| CORR2: BW - SW | LMM (oN) | 111 | 0.24 | .808 |
| CORR2: SL – SW | LMM (oN) | 129 | -0.25 | .806 |
| RAMP1: BL – BW | LMM (oN) | 119 | -0.09 | .932 |
| RAMP1: BL - SL | LMM (oN) | 136 | 0.20 | .845 |
| RAMP1: BW - SW | LMM (oN) | 111 | 0.76 | .452 |
| RAMP1: SL - SW | LMM (oN) | 129 | 0.43 | .666 |
| RAMP2: BL – BW | LMM (oN) | 119 | 0.23 | .817 |
| RAMP2: BL - SL | LMM (oN) | 126 | 1.64 | .104 |
| RAMP2: BW - SW | LMM (oN) | 111 | 0.65 | .515 |
| RAMP2: SL – SW | LMM (oN) | 119 | -0.81 | .421 |
| RAMP3: BL – BW | LMM (oN) | 119 | -0.80 | .427 |
| RAMP3: BL - SL | LMM (oN) | 119 | 0.65 | .515 |
| RAMP3: BW - SW | LMM (oN) | 111 | 1.03 | .304 |
| RAMP3: SL – SW | LMM (oN) | 111 | -0.46 | .649 |
| RAMP4: BL – BW | LMM (oN) | 126 | -0.05 | .962 |
| RAMP4: BL - SL | LMM (oN) | 119 | -0.20 | .839 |
| RAMP4: BW - SW | LMM (oN) | 119 | 0.61 | .540 |
| RAMP4: SL - SW | LMM (oN) | 111 | 0.79 | .433 |
| CIRC: BL – BW | LMM (oN) | 136 | 0.85 | .398 |
| CIRC: BL – SL | LMM (oN) | 126 | -0.09 | .931 |
| CIRC: BW – SW | LMM (oN) | 136 | 1.02 | .312 |
| CIRC: SL – SW | LMM (oN) | 126 | 2.00 | .048* |
| CENT: BL – BW | LMM (oN) | 111 | -0.37 | .713 |
| CENT: BL – SL | LMM (oN) | 111 | 1.97 | .052 |
| CENT: BW – SW | LMM (oN) | 111 | 2.49 | .014* |
| CENT: SL – SW | LMM (oN) | 111 | 0.16 | .876 |
| REST: BL – BW | LMM (oN) | 111 | -0.36 | .721 |
| REST: BL – SL | LMM (oN) | 111 | -0.59 | .559 |
| REST: BW - SW | LMM (oN) | 111 | 1.45 | .150 |
| REST: SL - SW | LMM (oN) | 111 | 1.68 | .096 |
| **zMCSF – Total duration in zone (zones)** | | | | |
| START: BL – BW | LMM (logit) | 265 | 0.52 | .603 |

| | | | | |
|---|---|---|---|---|
| START: BL - SL | LMM (logit) | 265 | -0.00 | .997 |
| START: BW - SW | LMM (logit) | 260 | -0.10 | .921 |
| START: SL – SW | LMM (logit) | 260 | 0.44 | .659 |
| DCR: BL – BW | LMM (logit) | 265 | -0.12 | .904 |
| DCR: BL - SL | LMM (logit) | 265 | -1.15 | .252 |
| DCR: BW - SW | LMM (logit) | 260 | -0.71 | .481 |
| DCR: SL – SW | LMM (logit) | 260 | 0.36 | .721 |
| CORR1: BL – BW | LMM (logit) | 260 | 0.55 | .581 |
| CORR1: BL - SL | LMM (logit) | 275 | 0.77 | .444 |
| CORR1: BW - SW | LMM (logit) | 260 | 1.38 | .169 |
| CORR1: SL – SW | LMM (logit) | 275 | 1.03 | .304 |
| CORN: BL – BW | LMM (logit) | 265 | -1.06 | .292 |
| CORN: BL - SL | LMM (logit) | 286 | 0.01 | .993 |
| CORN: BW - SW | LMM (logit) | 260 | 2.65 | .009** |
| CORN: SL – SW | LMM (logit) | 284 | 1.37 | .172 |
| CORR2: BL – BW | LMM (logit) | 265 | -0.98 | .327 |
| CORR2: BL - SL | LMM (logit) | 278 | 0.52 | .602 |
| CORR2: BW - SW | LMM (logit) | 260 | 2.27 | .024* |
| CORR2: SL – SW | LMM (logit) | 275 | 0.63 | .528 |
| RAMP1: BL – BW | LMM (logit) | 265 | 0.36 | .721 |
| RAMP1: BL - SL | LMM (logit) | 278 | 0.76 | .450 |
| RAMP1: BW - SW | LMM (logit) | 260 | 0.90 | .369 |
| RAMP1: SL - SW | LMM (logit) | 275 | 0.40 | .687 |
| RAMP2: BL – BW | LMM (logit) | 265 | 0.25 | .806 |
| RAMP2: BL - SL | LMM (logit) | 271 | 2.36 | .019* |
| RAMP2: BW - SW | LMM (logit) | 260 | 0.29 | .775 |
| RAMP2: SL – SW | LMM (logit) | 267 | -1.91 | .058 |
| RAMP3: BL – BW | LMM (logit) | 265 | 0.44 | .659 |
| RAMP3: BL - SL | LMM (logit) | 265 | 2.35 | .020* |
| RAMP3: BW - SW | LMM (logit) | 260 | 0.61 | .542 |
| RAMP3: SL – SW | LMM (logit) | 260 | -1.36 | .176 |
| RAMP4: BL – BW | LMM (logit) | 272 | -1.25 | .216 |
| RAMP4: BL - SL | LMM (logit) | 265 | 2.16 | .032* |
| RAMP4: BW - SW | LMM (logit) | 268 | 4.06 | < .001*** |
| RAMP4: SL - SW | LMM (logit) | 260 | 0.63 | .531 |
| CIRC: BL – BW | LMM (logit) | 279 | 0.31 | .761 |
| CIRC: BL – SL | LMM (logit) | 271 | 0.78 | .434 |
| CIRC: BW – SW | LMM (logit) | 281 | -0.02 | .984 |
| CIRC: SL – SW | LMM (logit) | 273 | -0.49 | .626 |
| CENT: BL – BW | LMM (logit) | 260 | 0.19 | .853 |
| CENT: BL – SL | LMM (logit) | 260 | 0.60 | .548 |
| CENT: BW – SW | LMM (logit) | 260 | -1.13 | .260 |
| CENT: SL – SW | LMM (logit) | 260 | -1.55 | .124 |
| REST: BL – BW | LMM (logit) | 260 | -0.17 | .867 |
| REST: BL – SL | LMM (logit) | 260 | 0.12 | .909 |
| REST: BW - SW | LMM (logit) | 260 | -0.22 | .824 |
| REST: SL - SW | LMM (logit) | 260 | -0.51 | .614 |
| **zMCSF – Duration of visit (zones)** | | | | |
| START: BL – BW | LMM (logit) | 247 | 0.28 | .784 |
| START: BL - SL | LMM (logit) | 247 | -1.95 | .053 |
| START: BW - SW | LMM (logit) | 241 | -0.49 | .627 |
| START: SL – SW | LMM (logit) | 241 | 1.81 | .072 |
| DCR: BL – BW | LMM (logit) | 247 | -0.27 | .789 |
| DCR: BL - SL | LMM (logit) | 247 | -1.63 | .104 |
| DCR: BW - SW | LMM (logit) | 241 | -0.27 | .785 |
| DCR: SL – SW | LMM (logit) | 241 | 1.14 | .257 |
| CORR1: BL – BW | LMM (logit) | 241 | 1.26 | .208 |
| CORR1: BL - SL | LMM (logit) | 259 | 0.00 | .997 |
| CORR1: BW - SW | LMM (logit) | 241 | 0.69 | .492 |
| CORR1: SL – SW | LMM (logit) | 259 | 1.82 | .070 |

| | | | | |
|---|---|---|---|---|
| CORN: BL – BW | LMM (logit) | 247 | -0.38 | .706 |
| CORN: BL - SL | LMM (logit) | 274 | 0.68 | .499 |
| CORN: BW - SW | LMM (logit) | 241 | 1.74 | .084 |
| CORN: SL – SW | LMM (logit) | 271 | 0.50 | .616 |
| CORR2: BL – BW | LMM (logit) | 247 | -0.78 | .438 |
| CORR2: BL - SL | LMM (logit) | 264 | 0.66 | .511 |
| CORR2: BW - SW | LMM (logit) | 241 | 0.44 | .658 |
| CORR2: SL – SW | LMM (logit) | 259 | -1.01 | .312 |
| RAMP1: BL – BW | LMM (logit) | 247 | 0.13 | .894 |
| RAMP1: BL - SL | LMM (logit) | 264 | 0.26 | .796 |
| RAMP1: BW - SW | LMM (logit) | 241 | 0.32 | .749 |
| RAMP1: SL - SW | LMM (logit) | 259 | 0.16 | .871 |
| RAMP2: BL – BW | LMM (logit) | 247 | -0.28 | .778 |
| RAMP2: BL - SL | LMM (logit) | 255 | 0.66 | .511 |
| RAMP2: BW - SW | LMM (logit) | 241 | -0.13 | .896 |
| RAMP2: SL – SW | LMM (logit) | 249 | -1.09 | .277 |
| RAMP3: BL – BW | LMM (logit) | 247 | 0.50 | .616 |
| RAMP3: BL - SL | LMM (logit) | 247 | 0.80 | .425 |
| RAMP3: BW - SW | LMM (logit) | 241 | -0.54 | .593 |
| RAMP3: SL – SW | LMM (logit) | 241 | -0.84 | .401 |
| RAMP4: BL – BW | LMM (logit) | 256 | -0.60 | .549 |
| RAMP4: BL - SL | LMM (logit) | 247 | 1.44 | .152 |
| RAMP4: BW - SW | LMM (logit) | 250 | 2.63 | .009** |
| RAMP4: SL - SW | LMM (logit) | 241 | 0.60 | .553 |
| CIRC: BL – BW | LMM (logit) | 265 | 0.33 | .739 |
| CIRC: BL – SL | LMM (logit) | 255 | 0.28 | .783 |
| CIRC: BW – SW | LMM (logit) | 267 | -0.18 | .862 |
| CIRC: SL – SW | LMM (logit) | 257 | -0.11 | .912 |
| CENT: BL – BW | LMM (logit) | 241 | 0.10 | .921 |
| CENT: BL – SL | LMM (logit) | 241 | -0.39 | .697 |
| CENT: BW – SW | LMM (logit) | 241 | -1.60 | .111 |
| CENT: SL – SW | LMM (logit) | 241 | -1.11 | .267 |
| REST: BL – BW | LMM (logit) | 241 | 0.39 | .694 |
| REST: BL – SL | LMM (logit) | 241 | 0.71 | .476 |
| REST: BW - SW | LMM (logit) | 241 | -0.79 | .431 |
| REST: SL - SW | LMM (logit) | 241 | -1.11 | .268 |
| **zMCSF – Number of visits to zone (zones)** | | | | |
| START: BL – BW | GLMM (NB) | | 0.21 | .830 |
| START: BL - SL | GLMM (NB) | | 1.12 | .263 |
| START: BW - SW | GLMM (NB) | | 0.01 | .989 |
| START: SL – SW | GLMM (NB) | | -0.92 | .357 |
| DCR: BL – BW | GLMM (NB) | | 0.05 | .964 |
| DCR: BL - SL | GLMM (NB) | | -0.30 | .761 |
| DCR: BW - SW | GLMM (NB) | | -0.55 | .581 |
| DCR: SL – SW | GLMM (NB) | | -0.19 | .852 |
| CORR1: BL – BW | GLMM (NB) | | -1.33 | .184 |
| CORR1: BL - SL | GLMM (NB) | | -0.11 | .917 |
| CORR1: BW - SW | GLMM (NB) | | 1.34 | .180 |
| CORR1: SL – SW | GLMM (NB) | | 0.12 | .906 |
| CORN: BL – BW | GLMM (NB) | | -0.78 | .436 |
| CORN: BL - SL | GLMM (NB) | | -0.10 | .924 |
| CORN: BW - SW | GLMM (NB) | | 1.87 | .061 |
| CORN: SL – SW | GLMM (NB) | | 1.09 | .277 |
| CORR2: BL – BW | GLMM (NB) | | -0.54 | .590 |
| CORR2: BL - SL | GLMM (NB) | | -0.67 | .502 |
| CORR2: BW - SW | GLMM (NB) | | 2.33 | .020* |
| CORR2: SL – SW | GLMM (NB) | | 2.37 | .018* |
| RAMP1: BL – BW | GLMM (NB) | | 0.18 | .855 |
| RAMP1: BL - SL | GLMM (NB) | | 0.22 | .824 |
| RAMP1: BW - SW | GLMM (NB) | | 0.97 | .333 |

| | | | | |
|---|---|---|---|---|
| RAMP1: SL - SW | GLMM (NB) | | 0.87 | .385 |
| RAMP2: BL – BW | GLMM (NB) | | 0.59 | .553 |
| RAMP2: BL - SL | GLMM (NB) | | 2.24 | .025* |
| RAMP2: BW - SW | GLMM (NB) | | 0.14 | .886 |
| RAMP2: SL – SW | GLMM (NB) | | -1.57 | .117 |
| RAMP3: BL – BW | GLMM (NB) | | 0.10 | .922 |
| RAMP3: BL - SL | GLMM (NB) | | 2.48 | .013* |
| RAMP3: BW - SW | GLMM (NB) | | 0.95 | .344 |
| RAMP3: SL – SW | GLMM (NB) | | -1.50 | .133 |
| RAMP4: BL – BW | GLMM (NB) | | -1.27 | .206 |
| RAMP4: BL - SL | GLMM (NB) | | 2.18 | .030* |
| RAMP4: BW - SW | GLMM (NB) | | 3.95 | < .001*** |
| RAMP4: SL - SW | GLMM (NB) | | 0.44 | .657 |
| CIRC: BL – BW | GLMM (NB) | | 0.40 | .688 |
| CIRC: BL – SL | GLMM (NB) | | 1.34 | .179 |
| CIRC: BW – SW | GLMM (NB) | | -0.27 | .786 |
| CIRC: SL – SW | GLMM (NB) | | -1.22 | .224 |
| CENT: BL – BW | GLMM (NB) | | 0.44 | .660 |
| CENT: BL – SL | GLMM (NB) | | 0.94 | .349 |
| CENT: BW – SW | GLMM (NB) | | -0.52 | .605 |
| CENT: SL – SW | GLMM (NB) | | -1.02 | .309 |
| REST: BL – BW | GLMM (NB) | | -0.10 | .923 |
| REST: BL – SL | GLMM (NB) | | -0.07 | .942 |
| REST: BW - SW | GLMM (NB) | | 0.20 | .841 |
| REST: SL - SW | GLMM (NB) | | 0.18 | .860 |
| **zMCSF – Number of visits (%) (zones)** | | | | |
| START: BL – BW | LMM (logit) | 306 | 0.33 | .742 |
| START: BL - SL | LMM (logit) | 306 | 0.54 | .590 |
| START: BW - SW | LMM (logit) | 306 | -0.30 | .763 |
| START: SL – SW | LMM (logit) | 306 | -0.52 | .605 |
| DCR: BL – BW | LMM (logit) | 306 | -0.19 | .852 |
| DCR: BL - SL | LMM (logit) | 306 | -0.93 | .352 |
| DCR: BW - SW | LMM (logit) | 306 | -0.92 | .357 |
| DCR: SL – SW | LMM (logit) | 306 | -0.15 | .880 |
| CORR1: BL – BW | LMM (logit) | 306 | -0.71 | .481 |
| CORR1: BL - SL | LMM (logit) | 308 | -0.10 | .918 |
| CORR1: BW - SW | LMM (logit) | 306 | 1.22 | .223 |
| CORR1: SL – SW | LMM (logit) | 308 | 0.58 | .563 |
| CORN: BL – BW | LMM (logit) | 306 | -1.81 | .072 |
| CORN: BL - SL | LMM (logit) | 310 | -1.38 | .168 |
| CORN: BW - SW | LMM (logit) | 306 | 2.48 | .014* |
| CORN: SL – SW | LMM (logit) | 309 | 1.95 | .052 |
| CORR2: BL – BW | LMM (logit) | 306 | -1.33 | .185 |
| CORR2: BL - SL | LMM (logit) | 308 | -0.73 | .469 |
| CORR2: BW - SW | LMM (logit) | 306 | 2.90 | .004** |
| CORR2: SL – SW | LMM (logit) | 308 | 2.16 | .031* |
| RAMP1: BL – BW | LMM (logit) | 306 | 0.10 | .917 |
| RAMP1: BL - SL | LMM (logit) | 308 | 0.28 | .783 |
| RAMP1: BW - SW | LMM (logit) | 306 | 0.71 | .478 |
| RAMP1: SL - SW | LMM (logit) | 308 | 0.47 | .636 |
| RAMP2: BL – BW | LMM (logit) | 306 | 0.26 | .797 |
| RAMP2: BL - SL | LMM (logit) | 307 | 2.32 | .021* |
| RAMP2: BW - SW | LMM (logit) | 306 | 0.19 | .851 |
| RAMP2: SL – SW | LMM (logit) | 307 | -1.95 | .052 |
| RAMP3: BL – BW | LMM (logit) | 306 | -0.04 | .966 |
| RAMP3: BL - SL | LMM (logit) | 306 | 2.09 | .037* |
| RAMP3: BW - SW | LMM (logit) | 306 | 1.00 | .316 |
| RAMP3: SL – SW | LMM (logit) | 306 | -1.20 | .229 |
| RAMP4: BL – BW | LMM (logit) | 308 | -1.76 | .080 |
| RAMP4: BL - SL | LMM (logit) | 306 | 1.29 | .200 |

| | | | | |
|---|---|---|---|---|
| RAMP4: BW - SW | LMM (logit) | 307 | 4.03 | < .001*** |
| RAMP4: SL - SW | LMM (logit) | 306 | 0.96 | .337 |
| CIRC: BL – BW | LMM (logit) | 309 | -0.01 | .994 |
| CIRC: BL – SL | LMM (logit) | 307 | -0.4 | .967 |
| CIRC: BW – SW | LMM (logit) | 309 | -0.47 | .641 |
| CIRC: SL – SW | LMM (logit) | 308 | -0.45 | .651 |
| CENT: BL – BW | LMM (logit) | 306 | 1.02 | .310 |
| CENT: BL – SL | LMM (logit) | 306 | 1.22 | .223 |
| CENT: BW – SW | LMM (logit) | 306 | -0.98 | .327 |
| CENT: SL – SW | LMM (logit) | 306 | -1.19 | .236 |
| REST: BL – BW | LMM (logit) | 306 | 0.19 | .847 |
| REST: BL – SL | LMM (logit) | 306 | -0.54 | .590 |
| REST: BW - SW | LMM (logit) | 306 | -0.27 | .787 |
| REST: SL - SW | LMM (logit) | 306 | 0.46 | .643 |
| **zMCSF – Latency to first entry (zones)** | | | | |
| START: BL – BW | LMM (oN) | 290 | 0.27 | .788 |
| START: BL - SL | LMM (oN) | 290 | 0.44 | .664 |
| START: BW - SW | LMM (oN) | 288 | 0.12 | .988 |
| START: SL – SW | LMM (oN) | 288 | -0.16 | .876 |
| DCR: BL – BW | LMM (oN) | 290 | 1.02 | .308 |
| DCR: BL - SL | LMM (oN) | 290 | 0.61 | .542 |
| DCR: BW - SW | LMM (oN) | 288 | -1.10 | .272 |
| DCR: SL – SW | LMM (oN) | 288 | -0.67 | .501 |
| CORR1: BL – BW | LMM (oN) | 288 | 0.80 | .426 |
| CORR1: BL - SL | LMM (oN) | 296 | 0.29 | .769 |
| CORR1: BW - SW | LMM (oN) | 288 | -2.28 | .024* |
| CORR1: SL – SW | LMM (oN) | 296 | -1.67 | .097 |
| CORN: BL – BW | LMM (oN) | 290 | 2.15 | .032* |
| CORN: BL - SL | LMM (oN) | 301 | 1.44 | .150 |
| CORN: BW - SW | LMM (oN) | 288 | -2.17 | .031* |
| CORN: SL – SW | LMM (oN) | 300 | -1.43 | .154 |
| CORR2: BL – BW | LMM (oN) | 290 | 0.12 | .903 |
| CORR2: BL - SL | LMM (oN) | 297 | -0.55 | .581 |
| CORR2: BW - SW | LMM (oN) | 288 | -2.08 | .039* |
| CORR2: SL – SW | LMM (oN) | 296 | -1.24 | .217 |
| RAMP1: BL – BW | LMM (oN) | 290 | -0.68 | .496 |
| RAMP1: BL - SL | LMM (oN) | 297 | -1.83 | .069 |
| RAMP1: BW - SW | LMM (oN) | 288 | -1.61 | .108 |
| RAMP1: SL - SW | LMM (oN) | 296 | -0.27 | .789 |
| RAMP2: BL – BW | LMM (oN) | 290 | -0.60 | .547 |
| RAMP2: BL - SL | LMM (oN) | 293 | -1.94 | .053 |
| RAMP2: BW - SW | LMM (oN) | 288 | -1.15 | .251 |
| RAMP2: SL – SW | LMM (oN) | 291 | 0.29 | .772 |
| RAMP3: BL – BW | LMM (oN) | 290 | -0.21 | .835 |
| RAMP3: BL - SL | LMM (oN) | 290 | -1.35 | .179 |
| RAMP3: BW - SW | LMM (oN) | 288 | -0.59 | .559 |
| RAMP3: SL – SW | LMM (oN) | 288 | 0.59 | .554 |
| RAMP4: BL – BW | LMM (oN) | 294 | -0.16 | .873 |
| RAMP4: BL - SL | LMM (oN) | 290 | -0.71 | .478 |
| RAMP4: BW - SW | LMM (oN) | 292 | -1.34 | .180 |
| RAMP4: SL - SW | LMM (oN) | 288 | -0.83 | .410 |
| CIRC: BL – BW | LMM (oN) | 298 | 1.95 | .052 |
| CIRC: BL – SL | LMM (oN) | 293 | 1.70 | .090 |
| CIRC: BW – SW | LMM (oN) | 299 | -1.73 | .084 |
| CIRC: SL – SW | LMM (oN) | 295 | -1.47 | .143 |
| CENT: BL – BW | LMM (oN) | 288 | -0.29 | .771 |
| CENT: BL – SL | LMM (oN) | 288 | -0.32 | .750 |
| CENT: BW – SW | LMM (oN) | 288 | -0.98 | .328 |
| CENT: SL – SW | LMM (oN) | 288 | -0.95 | .343 |
| REST: BL – BW | LMM (oN) | 288 | -0.29 | .774 |

| | | | | |
|---|---|---|---|---|
| REST: BL – SL | LMM (oN) | 288 | -0.01 | .989 |
| REST: BW - SW | LMM (oN) | 288 | 0.48 | .634 |
| REST: SL - SW | LMM (oN) | 288 | 0.20 | .839 |
