## Supplementary material for "Effects of stress coping styles and social defeat on zebrafish behaviour and brain transcriptomics": SF-3

**SF-3**: Statistical data of the significantly differentially expressed genes in bold and shy fish with different social experiences.

| **Ensembl** | **Gene** | **baseMean** | **LFC** | **LFC_SE** | $p_{val}$ | $p_{adj}$ |
|---|---|---|---|---|---|---|
| **HB: SL – SW (downregulated)** | | | | | | |
| ENSDARG00000103398 | fabp1b.2 | 9.97580 | -3.12e$^{-06}$ | .001443 | 3.15e$^{-06}$ | .0306 |
| ENSDARG00000103716 | si:busm1-194e12.11 | 39.9764 | -3.11e$^{-07}$ | .001443 | 6.93e$^{-14}$ | < .001 |
| ENSDARG00000092233 | vtg1 | 7.49148 | -6.82e$^{-07}$ | .001443 | 2.65e$^{-19}$ | < .001 |
| **HB: BL – BW (upregulated)** | | | | | | |
| ENSDARG00000114577 | si:dkey-159n16.2 | 41.6153 | 2.52316 | .646558 | 3.39e$^{-06}$ | .0036 |
| ENSDARG00000074547 | si:ch211-240l19.8 | 21.4076 | 4.31221 | 1.097784 | 2.87e$^{-06}$ | .0034 |
| ENSDARG00000038384 | th2 | 45.3254 | 2.58e$^{-06}$ | .001443 | 3.06e$^{-07}$ | .0010 |
| ENSDARG00000045139 | ca7 | 10.4512 | 4.38669 | 1.022823 | 8.26e$^{-07}$ | .0019 |
| ENSDARG00000090428 | ctrb1 | 63.6734 | 3.85813 | 1.090510 | 1.16e$^{-05}$ | .0093 |
| ENSDARG00000051923 | ccnb1 | 21.0396 | 2.15329 | .601519 | 1.26e$^{-05}$ | .0097 |
| ENSDARG00000007276 | ela3l | 30.7927 | 4.45723 | 1.178378 | 4.50e$^{-06}$ | .0044 |
| ENSDARG00000079497 | tcima | 1001.168 | .74411 | .194955 | 5.16e$^{-06}$ | .0058 |
| ENSDARG00000108939 | FQ378016.1 | 149.2342 | 5.12e$^{-06}$ | .001443 | 7.01e$^{-07}$ | .0019 |
| ENSDARG00000104773 | junbb | 1332.444 | 1.12732 | .231937 | 4.81e$^{-08}$ | < .001 |
| ENSDARG00000099195 | ier2a | 209.1977 | 1.20487 | .411377 | .00010 | .0487 |
| ENSDARG00000013856 | amy2a | 59.87223 | 4.38541 | 1.048393 | 8.65e$^{-07}$ | .0019 |
| ENSDARG00000075008 | pask | 27.61162 | 1.10010 | .370873 | .00010 | .0487 |
| ENSDARG00000060397 | hhip | 885.2998 | .55514 | .168520 | 3.81e$^{-05}$ | .0238 |
| ENSDARG00000074378 | junba | 202.3109 | .87789 | .224327 | 3.31e$^{-06}$ | .0036 |
| ENSDARG00000058471 | plk1 | 28.34337 | 1.71818 | .524452 | 3.64e$^{-05}$ | .0233 |
| ENSDARG00000061697 | ca14 | 8.674063 | 3.75428 | .883215 | 1.58e$^{-06}$ | .0025 |
| ENSDARG00000095796 | si:dkey-87o1.2 | 7.919607 | 2.87708 | .972471 | 9.84e$^{-05}$ | .0487 |
| ENSDARG00000079274 | prss59.1 | 89.94166 | 4.10973 | .995604 | 1.16e$^{-06}$ | .0022 |
| ENSDARG00000073742 | prss59.2 | 93.33689 | 4.06627 | .975937 | 9.67e$^{-07}$ | .0020 |
| ENSDARG00000031683 | fosab | 459.0697 | 1.65026 | .401785 | 1.51e$^{-06}$ | .0025 |
| ENSDARG00000104107 | nkx2.4b | 35.3621 | 1.46e$^{-06}$ | .001443 | 2.25e$^{-29}$ | < .001 |
| ENSDARG00000056765 | ela2l | 69.70546 | 3.71018 | 1.103459 | 2.12e$^{-05}$ | .0144 |
| ENSDARG00000056744 | ela2 | 34.00906 | 2.24264 | .772010 | .00011 | .0493 |
| ENSDARG00000100020 | pim1 | 1117.054 | .44426 | .099439 | 3.26e$^{-07}$ | .0010 |
| ENSDARG00000037421 | egr1 | 4246.108 | 1.39118 | .287517 | 5.37e$^{-08}$ | < .001 |
| ENSDARG00000055752 | npas4a | 2373.923 | 2.08054 | .419573 | 2.73e$^{-08}$ | < .001 |
| ENSDARG00000055250 | cntd2 | 30.65171 | 2.62734 | .769371 | 2.08e$^{-05}$ | .0143 |
| ENSDARG00000096645 | si:ch211-131k2.2 | 86.64902 | 4.42e$^{-06}$ | .001443 | 8.44e$^{-05}$ | .0467 |
| ENSDARG00000068846 | zgc:66024 | 9.001743 | 2.93778 | 1.00989 | .00011 | .0493 |
| ENSDARG00000000796 | nr4a1 | 137.5082 | 1.76833 | .475301 | 7.06e$^{-06}$ | .0061 |
| ENSDARG00000029822 | cel.2 | 12.40278 | 4.08896 | 1.03305 | 2.87e$^{-06}$ | .0034 |
| ENSDARG00000017490 | cel.1 | 23.16914 | 5.78811 | 1.47173 | 2.52e$^{-06}$ | .0034 |
| ENSDARG00000058682 | cd8b | 14.68529 | 3.44788 | .912050 | 5.62e$^{-06}$ | .0050 |
| ENSDARG00000056248 | si:dkey-183i3.5 | 34.77394 | 2.05416 | .576999 | 1.30e$^{-05}$ | .0097 |
| ENSDARG00000086881 | ier2b | 268.4214 | 1.26348 | .264806 | 7.61e$^{-08}$ | < .001 |
| ENSDARG00000094055 | si:dkey-88l16.3 | 10.07586 | 2.59750 | .851849 | 7.57e$^{-05}$ | .0435 |
| ENSDARG00000068976 | bsx | 44.76955 | 3.55e$^{-06}$ | .001443 | 1.71e$^{-06}$ | .0026 |
| ENSDARG00000045887 | mmp30 | 4.491145 | 2.03e$^{-06}$ | .001443 | 3.19e$^{-05}$ | .0211 |
| ENSDARG00000017314 | cela1.6 | 121.253 | 3.89265 | .989368 | 2.55e$^{-06}$ | .0034 |
| ENSDARG00000020298 | btg2 | 583.6895 | .81380 | .265443 | 6.86e$^{-05}$ | .0405 |
| ENSDARG00000097513 | CT573383.1 | 35.69544 | 4.00335 | 1.39249 | 9.25e$^{-05}$ | .0487 |
| ENSDARG00000045638 | slc13a1 | 73.23511 | 1.59e$^{-06}$ | .001443 | 1.82e$^{-05}$ | .0132 |
| ENSDARG00000061416 | c2cd4a | 36.23096 | 1.28e$^{-05}$ | .001443 | .00010 | .0487 |
| ENSDARG00000098739 | h2af1al | 45.33396 | 2.21960 | .760912 | .00010 | .0487 |
| **HB: BL – BW (downregulated)** | | | | | | |
| ENSDARG00000098462 | CU570782.1 | 10.2594 | -2.29e$^{-06}$ | .001443 | 6.55e$^{-05}$ | .0397 |
| ENSDARG00000001870 | atp1a1a.4 | 13.30379 | -3.30e$^{-06}$ | .001443 | 8.53e$^{-05}$ | .0467 |
| ENSDARG00000096273 | si:dkey-3n22.9 | 63.31436 | -8.69e$^{-07}$ | .001443 | 4.37e$^{-06}$ | .0044 |
| ENSDARG00000094550 | BX649490.2 | 29.53691 | -5.68e$^{-08}$ | .001443 | 9.20e$^{-06}$ | .0076 |

| | | | | | | |
|---|---|---|---|---|---|---|
| ENSDARG00000111856 | BX649490.4 | 27.61877 | -6.92e-08 | .001443 | 9.45e-05 | .0487 |
| **HB: SL – BL (upregulated)** | | | | | | |
| ENSDARG00000096187 | si:dkey-21h14.10 | 4.678902 | .00392 | .041370 | 8.92e-05 | .0256 |
| ENSDARG00000070314 | cald1a | 255.1406 | .44331 | .127597 | 2.07e-05 | .0102 |
| ENSDARG00000106267 | CR388042.1 | 35.03077 | 5.24287 | .869431 | 9.57e-11 | < .001 |
| ENSDARG00000035820 | drd4b | 8.123519 | .01271 | .044010 | 2.99e-05 | .0128 |
| ENSDARG00000100738 | osmr | 38.80564 | .00421 | .041412 | 7.94e-05 | .0243 |
| ENSDARG00000074320 | KCNAB3 | 228.6036 | .51040 | .179798 | .00014 | .0338 |
| ENSDARG00000100972 | myh11b | 5.909423 | 4.59848 | 1.304109 | 2.06e-05 | .0101 |
| ENSDARG00000098749 | tbcd | 553.2857 | .29519 | .104841 | .00018 | .0398 |
| ENSDARG00000021720 | col7a1 | 48.52985 | 1.66307 | .474471 | 1.64e-05 | .0092 |
| ENSDARG00000001913 | palmda | 74.66797 | .002508 | .041210 | 4.32e-05 | .0159 |
| ENSDARG00000016994 | ssrp1b | 983.9128 | .33829 | .081903 | 1.83e-06 | .0024 |
| ENSDARG00000061629 | ndufaf5 | 240.9738 | .29807 | .101821 | .00011 | .0298 |
| ENSDARG00000096586 | si:ch1073-110a20.3 | 7.755719 | 4.51644 | 1.237995 | 1.21e-05 | .0083 |
| ENSDARG00000020028 | cps1 | 11.81904 | 1.91559 | .594821 | 4.73e-05 | .0164 |
| ENSDARG00000102888 | gpr39 | 12.16084 | .00786 | .042198 | 4.66e-05 | .0164 |
| ENSDARG00000032838 | si:dkey-206f10.1 | 3.230056 | 6.33595 | 2.637496 | 9.21e-05 | .0261 |
| ENSDARG00000098779 | BX547930.4 | 7.434990 | .00233 | .041194 | 8.25e-05 | .0249 |
| ENSDARG00000069407 | zgc:194990 | 109.3109 | .71567 | .261430 | .00019 | .0415 |
| ENSDARG00000036156 | fnbp1a | 283.9691 | .59804 | .217282 | .00019 | .0413 |
| ENSDARG00000088116 | gstm.3 | 34.85850 | 2.24500 | .612938 | 8.76e-06 | .0688 |
| ENSDARG00000062487 | si:dkey-6n6.1 | 78.40270 | 1.66594 | .308191 | 2.76e-09 | < .001 |
| ENSDARG00000110838 | agbl4 | 75.61300 | .60852 | .176271 | 2.19e-05 | .0103 |
| ENSDARG00000116896 | TMEM233 | 9.426251 | 3.69603 | .928925 | 4.48e-06 | .0044 |
| ENSDARG00000022503 | pkd2l1 | 12.70354 | 2.59294 | .970767 | .000216 | .0420 |
| ENSDARG00000058486 | caps2 | 54.22949 | .84579 | .296854 | .000161 | .0365 |
| ENSDARG00000010296 | kcnh6b | 16.03450 | 3.9770 | 1.097234 | 9.23e-06 | .0070 |
| ENSDARG00000016088 | rtn2a | 60.25248 | .84956 | .236418 | 1.28e-05 | .0084 |
| ENSDARG00000112977 | CABZ01061591.1 | 187.6869 | .42882 | .162103 | .00027 | .0474 |
| ENSDARG00000059202 | tspan2b | 69.85738 | 1.06775 | .370554 | .00013 | .0317 |
| ENSDARG00000029905 | phyhd1 | 133.7733 | .81975 | .231780 | 1.50e-05 | .0088 |
| ENSDARG00000005141 | camkvb | 2239.715 | .59149 | .220080 | .00023 | .0437 |
| ENSDARG00000074808 | megf6b | 45.56442 | 1.33145 | .490102 | .00020 | .0418 |
| ENSDARG00000089645 | si:ch1073-406l10.2 | 71.83379 | .70730 | .248980 | .00016 | .0360 |
| ENSDARG00000008790 | actr3b | 300.7535 | .69241 | .207170 | 2.59e-05 | .0114 |
| ENSDARG00000026165 | col11a1a | 78.69788 | .90149 | .333775 | .00022 | .0420 |
| ENSDARG00000071685 | slco5a1a | 169.7608 | .47029 | .173953 | .00022 | .0420 |
| ENSDARG00000090145 | tmem240b | 367.1843 | .53664 | .176716 | 8.47e-05 | .0250 |
| ENSDARG00000094550 | BX649490.2 | 29.53691 | .00038 | .041099 | 2.90e-06 | .0030 |
| ENSDARG00000095798 | BX666064.1 | 24.36383 | 2.88525 | .998531 | .00011 | .0295 |
| ENSDARG00000063180 | dock3 | 1869.461 | .53576 | .183241 | .00012 | .0298 |
| ENSDARG00000111856 | BX649490.4 | 27.61877 | .00039 | .041099 | 2.14e-06 | .0026 |
| ENSDARG00000071076 | ldhbb | 839.0020 | .62288 | .189858 | 3.84e-05 | .0148 |
| ENSDARG00000104235 | myo5c | 103.0197 | 2.25147 | .624744 | 1.09e-05 | .0077 |
| ENSDARG00000032650 | fuk | 194.8692 | .51119 | .147328 | 2.03e-05 | .0102 |
| ENSDARG00000103230 | CABZ01079302.1 | 24.87903 | 3.54540 | .766015 | 1.55e-07 | < .001 |
| **HB: SL – BL (downregulated)** | | | | | | |
| ENSDARG00000022165 | mgst1.2 | 284.038 | -.93278 | .319116 | .00011 | .0295 |
| ENSDARG00000038384 | th2 | 45.3253 | -6.41714 | 1.704624 | 4.54e-08 | < .001 |
| ENSDARG00000017010 | scamp2l | 18.77712 | -.02040 | .048802 | .00020 | .0418 |
| ENSDARG00000088087 | kdm6ba | 1696.720 | -.46403 | .135927 | 2.61e-05 | .0114 |
| ENSDARG00000010332 | zgc:56231 | 11.37156 | -.00536 | .041610 | 3.51e-05 | .0144 |
| ENSDARG00000051923 | ccnb1 | 21.03961 | -.01098 | .043257 | 3.13e-05 | .0132 |
| ENSDARG00000089769 | hapln1a | 44.89787 | -.01203 | .043686 | .00014 | .0333 |
| ENSDARG00000035018 | thy1 | 82.86500 | -.01171 | .043552 | .00011 | .0295 |
| ENSDARG00000005943 | htra4 | 15.05444 | -.00273 | 0,041232 | 1.24e-05 | .0083 |
| ENSDARG00000079497 | tcima | 1001.168 | -.63156 | 0,199658 | 5.77e-05 | .0189 |
| ENSDARG00000108939 | FQ378016.1 | 149.2342 | -.00813 | 0,042288 | 5.52e-06 | .0048 |
| ENSDARG00000070484 | zgc:195001 | 58.24178 | -.00812 | 0,042268 | .00012 | .0317 |

| | | | | | | |
|---|---|---|---|---|---|---|
| ENSDARG00000038133 | zgc:113411 | 78.79128 | -1.00418 | 0,333284 | $8.59e^{-05}$ | .0250 |
| ENSDARG00000037782 | sox8b | 47.35481 | -.02126 | 0,049479 | .00026 | .0464 |
| ENSDARG00000087012 | BX004816.2 | 18.04599 | -.00669 | 0,041896 | $4.51e^{-05}$ | .0161 |
| ENSDARG00000044511 | etv5b | 1142.309 | -.94065 | 0,275892 | $2.24e^{-05}$ | .0103 |
| ENSDARG00000055192 | zgc:136930 | 7.717208 | -.00252 | 0,041211 | .00020 | .0415 |
| ENSDARG00000033231 | mcm6l | 8.914312 | -.00574 | 0,041685 | $4.17e^{-05}$ | .0156 |
| ENSDARG00000097559 | cyp8b3 | 15.43430 | -.00488 | 0,041525 | $2.77e^{-06}$ | .0030 |
| ENSDARG00000100741 | cdc20 | 30.96875 | -.01199 | 0,043652 | .00026 | .0464 |
| ENSDARG00000094449 | BX511231.1 | 14.95514 | -.00200 | 0,041168 | .00014 | .0338 |
| ENSDARG00000002330 | lhx8a | 48.80534 | -.00516 | 0,041564 | .00022 | .0420 |
| ENSDARG00000076043 | si:dkeyp-73d8.9 | 38.31173 | -.00975 | 0,042798 | $5.21e^{-05}$ | .0178 |
| ENSDARG00000069018 | cyp7a1 | 9.777166 | -.00379 | 0,041355 | $3.69e^{-05}$ | .0147 |
| ENSDARG00000104907 | ing2 | 215.8740 | -.36566 | 0,124877 | .00013 | .0321 |
| ENSDARG00000058471 | plk1 | 28.34337 | -2.32654 | 0,503943 | $1.50e^{-07}$ | .0003 |
| ENSDARG00000075929 | si:dkey-219c10.4 | 35.78369 | -2.70998 | 0,807650 | $3.55e^{-05}$ | .0144 |
| ENSDARG00000069763 | etv5a | 2830.509 | -0.74446 | 0,217520 | $2.20e^{-05}$ | .0103 |
| ENSDARG00000079964 | dlx2a | 105.7694 | -2.32205 | 0,624742 | $1.48e^{-05}$ | .0088 |
| ENSDARG00000061697 | ca14 | 8.674063 | -.00865 | 0,042429 | $7.56e^{-05}$ | .0234 |
| ENSDARG00000093760 | si:ch211-197h24.9 | 69.90477 | -.02474 | 0,052808 | .00016 | .0365 |
| ENSDARG00000043457 | gapdh | 393.0814 | -.00875 | 0,042474 | $1.44e^{-05}$ | .0088 |
| ENSDARG00000029259 | zgc:136493 | 213.7953 | -1.33699 | 0,262303 | $1.32e^{-08}$ | < .001 |
| ENSDARG00000070656 | si:ch211-69g19.2 | 44.87583 | -.01210 | 0,043713 | .00013 | .0329 |
| ENSDARG00000060862 | atxn1b | 2255.713 | -.39883 | 0,152312 | .00023 | .0442 |
| ENSDARG00000015543 | s100a1 | 19.60989 | -.00788 | 0,042205 | $1.76e^{-05}$ | .0096 |
| ENSDARG00000021787 | abcb5 | 20.48300 | -.00445 | 0,041451 | $6.83e^{-05}$ | .0214 |
| ENSDARG00000014039 | si:dkeyp-93d12.1 | 6.454126 | -.00871 | 0,042439 | .00017 | .0382 |
| ENSDARG00000088989 | si:dkey-241l7.5 | 60.03630 | -.00269 | 0,041229 | $1.06e^{-09}$ | < .001 |
| ENSDARG00000105335 | si:dkey-241l7.2 | 34.39383 | -.00143 | 0,041134 | $6.70e^{-07}$ | .0012 |
| ENSDARG00000094752 | rpe65b | 66.73917 | -1.09521 | 0,262732 | $1.30e^{-06}$ | .0019 |
| ENSDARG00000002295 | si:dkey-21p1.3 | 6.842278 | -.00656 | 0,041858 | $6.16e^{-05}$ | .0196 |
| ENSDARG00000010572 | slc25a25a | 79.52563 | -.02183 | 0,050016 | .00017 | .0387 |
| ENSDARG00000057433 | st6galnac5b | 168.9028 | -.61126 | 0,177332 | $2.05e^{-05}$ | .0102 |
| ENSDARG00000058865 | endog | 119.0462 | -.60587 | 0,177204 | $2.40e^{-05}$ | .0109 |
| ENSDARG00000078216 | eps15 | 887.4424 | -.42562 | 0,130966 | $4.43e^{-05}$ | .0160 |
| ENSDARG00000100020 | pim1 | 1117.054 | -.52578 | 0,097541 | $2.50e^{-09}$ | < .001 |
| ENSDARG00000103716 | si:busm1-194e12.11 | 39.97644 | -.00051 | 0,041101 | $3.82e^{-13}$ | < .001 |
| ENSDARG00000015657 | zgc:77112 | 39.23900 | -.01116 | 0,043357 | $4.76e^{-06}$ | .0045 |
| ENSDARG00000071021 | papss2a | 68.39562 | -.01951 | 0,048238 | $1.40e^{-05}$ | .0088 |
| ENSDARG00000040623 | fosl2 | 315.2186 | -.02192 | 0,050121 | .00016 | .0360 |
| ENSDARG00000100558 | slbp | 101.0754 | -.74040 | 0,236396 | $5.93e^{-05}$ | .0191 |
| ENSDARG00000075421 | pttg1 | 9.603928 | -.00721 | 0,042016 | .00024 | .0448 |
| ENSDARG00000037393 | slc43a1a | 7.290583 | -.00595 | 0,041720 | .00026 | .0464 |
| ENSDARG00000055752 | npas4a | 2373.923 | -.33653 | 0,446595 | $8.48e^{-05}$ | .0250 |
| ENSDARG00000002197 | pygl | 194.4050 | -.02214 | 0,050379 | $5.39e^{-05}$ | .0181 |
| ENSDARG00000055250 | cntd2 | 30.65171 | -.00789 | 0,042212 | $1.60e^{-05}$ | .0092 |
| ENSDARG00000076659 | cdca7b | 15.20359 | -.01268 | 0,043955 | .00028 | .0489 |
| ENSDARG00000068846 | zgc:66024 | 9.001743 | -.00592 | 0,041728 | $6.11e^{-06}$ | .0051 |
| ENSDARG00000039007 | eno3 | 573.5022 | -.05443 | 0,152594 | .00011 | .0295 |
| ENSDARG00000000796 | nr4a1 | 137.5082 | -.01354 | 0,044377 | .00024 | .0448 |
| ENSDARG00000006603 | csrp1a | 66.00428 | -.54570 | 0,202350 | .00024 | .0443 |
| ENSDARG00000054211 | st8sia7.1 | 2.054069 | -.00378 | 0,041353 | $2.00e^{-05}$ | .0102 |
| ENSDARG00000101595 | tgm1l2 | 24.06534 | -.00198 | 0,041167 | .00028 | .0492 |
| ENSDARG00000053395 | cdkn2aipnl | 336.8879 | -.35778 | 0,132669 | .00021 | .0420 |
| ENSDARG00000058682 | cd8b | 14.68529 | -.00663 | 0,041890 | $6.42e^{-07}$ | .0012 |
| ENSDARG00000044691 | ppp1r3b | 10.14181 | -.00671 | 0,041895 | .00014 | .0338 |
| ENSDARG00000025174 | zgc:103482 | 32.46348 | -.01382 | 0,044605 | $1.17e^{-06}$ | .0018 |
| ENSDARG00000018757 | klf5l | 3.058990 | -.00401 | 0,041381 | .00018 | .0394 |
| ENSDARG00000117089 | CKS2 | 21.87011 | -.01652 | 0,046098 | $3.82e^{-05}$ | .0148 |
| ENSDARG00000068732 | spry4 | 443.2542 | -.53375 | 0,204834 | .00027 | .0480 |
| ENSDARG00000086881 | ier2b | 268.4214 | -1.11077 | 0,267289 | $1.09e^{-06}$ | .0018 |

| | | | | | | |
|---|---|---|---|---|---|---|
| ENSDARG00000089802 | akap1a | 2.358948 | -.00400 | 0,041384 | $4.99e^{-06}$ | .0045 |
| ENSDARG00000068976 | bsx | 44.76955 | -.00534 | 0,041601 | .00021 | .0420 |
| ENSDARG00000071735 | prlh2 | 46.78754 | -.00494 | 0,041535 | $2.47e^{-06}$ | .0028 |
| ENSDARG00000045306 | slc51a | 37.97154 | -.01404 | 0,044667 | .00011 | .0295 |
| ENSDARG00000045453 | f13a1a.1 | 49.04331 | -.00343 | 0,041304 | .00022 | .0427 |
| ENSDARG00000038429 | csrnp1b | 474.4105 | -.52803 | 0,166938 | $5.56e^{-05}$ | .0184 |
| ENSDARG00000071662 | si:rp71-36a1.3 | 27.97897 | -.00121 | 0,041123 | .00010 | .0287 |
| ENSDARG00000092233 | vtg1 | 7.491476 | -.00193 | 0,041164 | $2.21e^{-37}$ | < .001 |
| ENSDARG00000022372 | kng1 | 34.04275 | -4.29655 | 0,606940 | $9.91e^{-14}$ | < .001 |
| ENSDARG00000102493 | ticam1 | 145.5738 | -.67717 | 0,208460 | $4.02e^{-05}$ | .0153 |
| ENSDARG00000002365 | cers5 | 209.1414 | -.59727 | 0,164092 | $1.02e^{-05}$ | .0075 |
| ENSDARG00000053918 | srfa | 221.9119 | -.46776 | 0,171538 | .00021 | .0420 |
| ENSDARG00000103183 | CABZ01034691.1 | 10.06589 | -.00665 | 0,041896 | $1.39e^{-06}$ | .0019 |
| ENSDARG00000079645 | sc:d217 | 179.2094 | -.01378 | 0,044561 | $7.46e^{-06}$ | .0061 |
| ENSDARG00000114409 | CR450686.6 | 33.19857 | -.00676 | 0,041906 | .00015 | .0360 |
| ENSDARG00000102414 | myhz1.1 | 100.4280 | -.00362 | 0,041334 | $2.45e^{-06}$ | .0028 |
| ENSDARG00000098360 | cyp19a1b | 10665.86 | -.00608 | 0,041749 | .00027 | .0474 |
| ENSDARG00000101164 | nansb | 1.959533 | -.00672 | 0,041894 | .00021 | .0420 |
| ENSDARG00000104040 | CABZ01046088.1 | 7.020808 | -.01344 | 0,044325 | .00020 | .0418 |
| **HB: SW – BW (upregulated)** | | | | | | |
| ENSDARG00000088836 | si:ch211-76m11.5 | 24.21246 | .00059 | .0115717 | $6.47e^{-05}$ | .0445 |
| ENSDARG00000106267 | CR388042.1 | 35.03077 | 6.16282 | .9884124 | $6.45e^{-11}$ | < .001 |
| ENSDARG00000114783 | znf985 | 4.228667 | .00045 | .0115626 | $3.62e^{-05}$ | .0281 |
| ENSDARG00000028912 | si:dkey-10h3.2 | 14.98272 | 3.21717 | .7314620 | $4.52e^{-07}$ | .0011 |
| ENSDARG00000018077 | rbp1.1 | 114.7749 | 1.63333 | .3958970 | $1.34e^{-06}$ | .0029 |
| ENSDARG00000014790 | g3bp2 | 1418.021 | 1.26125 | .3527051 | $1.12e^{-05}$ | .0123 |
| ENSDARG00000102888 | gpr39 | 12.16084 | 4.37053 | .9151636 | $1.08e^{-07}$ | < .001 |
| ENSDARG00000104107 | nkx2.4b | 35.36210 | .00017 | .0115517 | $7.92e^{-26}$ | < .001 |
| ENSDARG00000075405 | adck5 | 181.5564 | .72712 | .1535012 | $8.49e^{-08}$ | < .001 |
| ENSDARG00000088898 | caln1 | 443.8570 | .61575 | .2014710 | $7.45e^{-05}$ | .0460 |
| ENSDARG00000093931 | rflnb | 350.3003 | .47358 | .1490874 | $5.23e^{-05}$ | .0382 |
| ENSDARG00000102050 | MCOLN3 | 22.21918 | 2.74090 | .8918445 | $6.30e^{-05}$ | .0445 |
| ENSDARG00000020811 | efemp2b | 225.5847 | .87753 | .2566612 | $2.25e^{-05}$ | .0193 |
| ENSDARG00000019713 | oatx | 39.21077 | .00021 | .0115528 | $1.59e^{-05}$ | .0153 |
| ENSDARG00000023609 | AL845324.1 | 1102.719 | .49986 | .1343324 | $7.32e^{-06}$ | .0099 |
| ENSDARG00000074808 | megf6b | 45.56442 | 1.70800 | .4704327 | $1.04e^{-05}$ | .0123 |
| ENSDARG00000094550 | BX649490.2 | 29.53691 | $2.95e^{-05}$ | .0115500 | $5.95e^{-06}$ | .0099 |
| ENSDARG00000111856 | BX649490.4 | 27.61877 | $2.96e^{-05}$ | .0115500 | $4.36e^{-06}$ | .0087 |
| ENSDARG00000045638 | slc13a1 | 73.23511 | .00021 | .0115528 | $6.66e^{-08}$ | < .001 |
| ENSDARG00000100332 | CABZ01084942.1 | 12.94216 | .00043 | .0115614 | $2.32e^{-05}$ | .0193 |
| ENSDARG00000100540 | CABZ01114105.1 | 47.84465 | 2.82088 | .7682446 | $8.09e^{-06}$ | .0103 |
| **HB: SW – BW (downregulated)** | | | | | | |
| ENSDARG00000098932 | gigyf1a | 514.4630 | -.53129 | .172418 | $6.84e^{-05}$ | .0448 |
| ENSDARG00000023495 | ift74 | 525.1150 | -.00483 | .013054 | $7.07e^{-05}$ | .0448 |
| ENSDARG00000037805 | lgals3bpa | 55.73727 | -.00069 | .011580 | $2.00e^{-05}$ | .0179 |
| ENSDARG00000037782 | sox8b | 47.35481 | -.00181 | .011754 | $6.53e^{-06}$ | .0099 |
| ENSDARG00000055589 | s100t | 3.372115 | -.00028 | .011555 | $6.94e^{-06}$ | .0099 |
| ENSDARG00000029259 | zgc:136493 | 213.7953 | -.00305 | .012150 | $7.20e^{-10}$ | < .001 |
| ENSDARG00000010478 | hsp90aa1.1 | 50.37898 | -.00083 | .011593 | $7.29e^{-06}$ | .0099 |
| ENSDARG00000088989 | si:dkey-241l7.5 | 60.03630 | -.00020 | .011553 | $7.93e^{-05}$ | .0478 |
| ENSDARG00000105214 | agtpbp1 | 493.3345 | -.30493 | .100181 | $6.99e^{-05}$ | .0448 |
| ENSDARG00000053481 | entpd5a | 25.66867 | -.00087 | .011597 | $1.76e^{-05}$ | .0163 |
| ENSDARG00000031483 | col9a1b | 11.06776 | -.00102 | .011615 | $4.93e^{-05}$ | .0371 |
| ENSDARG00000096327 | cd164l2 | 14.68670 | -.00098 | .011611 | $2.46e^{-07}$ | < .001 |
| ENSDARG00000096273 | si:dkey-3n22.9 | 63.31436 | -.00011 | .011551 | $7.38e^{-06}$ | .0099 |
| ENSDARG00000103586 | si:dkey-65j6.2 | 22.36293 | -.00141 | .011674 | $1.36e^{-05}$ | .0142 |
| ENSDARG00000054319 | oxct1b | 16.58725 | -.00045 | .011563 | $8.13e^{-05}$ | .0478 |
| ENSDARG00000087633 | si:dkey-11o18.5 | 118.2286 | -.00040 | .011560 | $2.57e^{-08}$ | < .001 |
| ENSDARG00000089357 | miga2 | 164.2681 | -.00418 | .012673 | $1.42e^{-05}$ | .0142 |

| | | | | | | |
|---|---|---|---|---|---|---|
| ENSDARG00000105511 | BX248521.2 | 38.42651 | -1.39769 | .388132 | $1.10e^{-05}$ | .0123 |
| ENSDARG00000022372 | kng1 | 34.04275 | -4.54008 | .631777 | $5.27e^{-14}$ | < .001 |
| ENSDARG00000104467 | bglapl | 16.29623 | -.00086 | .011595 | $2.80e^{-05}$ | .0225 |
| **FBMB: SL – SW (downregulated)** | | | | | | |
| ENSDARG00000103716 | si:busm1-194e12.11 | 39.97644 | $-3.13e^{-07}$ | .001443 | $3.12e^{-12}$ | $9.11e^{-08}$ |
| **FBMB: BL – BW (upregulated)** | | | | | | |
| ENSDARG00000096243 | ighv1-3 | 5.821612 | $3.66e^{-07}$ | .001443 | $3.89e^{-09}$ | < .001 |
| ENSDARG00000061697 | ca14 | 8.674063 | 3.04691 | .763709 | $2.84e^{-06}$ | .0277 |
| ENSDARG00000002347 | cyp11a1 | 11.00258 | $2.80e^{-07}$ | .001443 | $9.45e^{-08}$ | .0014 |
| **FBMB: SL – BL (upregulated)** | | | | | | |
| ENSDARG00000106267 | CR388042.1 | 35.03077 | 4.09343 | .783176 | $1.03e^{-08}$ | < .001 |
| ENSDARG00000100738 | osmr | 38.80564 | 3.80891 | 1.187676 | $3.53e^{-05}$ | .0290 |
| ENSDARG00000020822 | ift22 | 281.2796 | .51977 | .172667 | $9.10e^{-05}$ | .0498 |
| ENSDARG00000097774 | CR387996.1 | 3.468637 | .00121 | .019703 | $7.24e^{-05}$ | .0450 |
| ENSDARG00000102888 | gpr39 | 12.16084 | .27457 | .900024 | $9.38e^{-06}$ | .0135 |
| ENSDARG00000007783 | blk | 53.89848 | .25118 | .663937 | $2.32e^{-05}$ | .0205 |
| ENSDARG00000062487 | si:dkey-6n6.1 | 78.40270 | 1.28260 | .320015 | $2.36e^{-06}$ | .0068 |
| ENSDARG00000076994 | adgra2 | 29.16110 | .97243 | .293805 | $3.76e^{-05}$ | .0298 |
| ENSDARG00000116896 | TMEM233 | 9.426251 | 3.28898 | .889375 | $9.09e^{-06}$ | .0135 |
| ENSDARG00000010296 | kcnh6b | 16.03450 | 4.26442 | 1.242354 | $2.03e^{-05}$ | .0203 |
| ENSDARG00000029905 | phyhd1 | 133.7733 | .78310 | .228669 | $2.21e^{-05}$ | .0203 |
| ENSDARG00000074808 | megf6b | 45.56442 | 1.72066 | .475471 | $1.01e^{-05}$ | .0135 |
| ENSDARG00000090219 | wdr45 | 561.4647 | .23736 | .076515 | $6.75e^{-05}$ | .0431 |
| ENSDARG00000008790 | actr3b | 300.7535 | .64454 | .215436 | $9.01e^{-05}$ | .0498 |
| ENSDARG00000042753 | cts12 | 49.14525 | .93903 | .308335 | $7.73e^{-05}$ | .0466 |
| ENSDARG00000094550 | BX649490.2 | 29.53691 | $8.60e^{-05}$ | .019649 | $5.11e^{-06}$ | .0098 |
| ENSDARG00000104015 | fgfr1bl | 21.49476 | 2.36559 | .573686 | $1.54e^{-06}$ | .0051 |
| ENSDARG00000071339 | borcs8 | 139.5423 | .45237 | .124500 | $1.06e^{-05}$ | .0135 |
| ENSDARG00000111856 | BX649490.4 | 27.61877 | $8.60e^{-05}$ | .019649 | $4.77e^{-06}$ | .0098 |
| ENSDARG00000071076 | ldhbb | 839.0020 | .77107 | .186599 | $1.44e^{-06}$ | .0051 |
| ENSDARG00000104235 | myo5c | 103.0197 | 2.39394 | .622511 | $4.31e^{-06}$ | .0098 |
| ENSDARG00000103230 | CABZ01079302.1 | 24.87903 | 3.68635 | .810422 | $2.19e^{-07}$ | .0013 |
| ENSDARG00000100284 | cd247 | 49.10169 | 1.32348 | .372279 | $1.37e^{-05}$ | .0166 |
| **FBMB: SL – BL (downregulated)** | | | | | | |
| ENSDARG00000037805 | lgals3bpa | 55.73727 | -.00190 | .019782 | $8.55e^{-05}$ | .0491 |
| ENSDARG00000037782 | sox8b | 47.35481 | -.00488 | .020532 | $4.99e^{-05}$ | .0359 |
| ENSDARG00000075785 | si:ch73-190m4.1 | 18.32180 | -.00238 | .019859 | $1.70e^{-05}$ | .0177 |
| ENSDARG00000113332 | CABZ01084501.2 | 54.06207 | -.00270 | .019918 | $4.43e^{-05}$ | .0340 |
| ENSDARG00000029259 | zgc:136493 | 213.7953 | -1.02621 | .275132 | $7.02e^{-06}$ | .0115 |
| ENSDARG00000070604 | zgc:162509 | 10.21273 | -2.28453 | .640860 | $1.61e^{-05}$ | .0176 |
| ENSDARG00000057433 | st6galnac5b | 168.9028 | -.01041 | .023933 | $2.18e^{-05}$ | .0203 |
| ENSDARG00000103716 | si:busm1-194e12.11 | 39.97644 | -.00012 | .019650 | $5.89e^{-13}$ | < .001 |
| ENSDARG00000105392 | si:ch73-22o18.1 | 14.36229 | -.00109 | .019693 | $4.95e^{-05}$ | .0359 |
| ENSDARG00000037403 | hspa8b | 2744.924 | -.83804 | .269041 | $6.14e^{-05}$ | .0415 |
| ENSDARG00000037781 | acss2 | 700.6046 | -.21979 | .062591 | $1.55e^{-05}$ | .0176 |
| ENSDARG00000068732 | spry4 | 443.2542 | -.61872 | .197473 | $5.29e^{-05}$ | .0368 |
| ENSDARG00000002494 | itgb6 | 42.03611 | -.00160 | .019744 | $4.65e^{-06}$ | .0098 |
| ENSDARG00000075192 | yme1l1a | 370.0990 | -.64764 | .138760 | $1.25e^{-07}$ | < .001 |
| ENSDARG00000022372 | kng1 | 34.04275 | -.00308 | .020004 | $4.06e^{-07}$ | .0019 |
| ENSDARG00000103183 | CABZ01034691.1 | 10.06589 | -3.5565 | 1.053463 | $3.22e^{-05}$ | .0274 |
| ENSDARG00000079645 | sc:d217 | 179.2094 | -.00309 | .020004 | $5.74e^{-06}$ | .0101 |
| ENSDARG00000077357 | lrrc61 | 129.8692 | -.00868 | .022533 | $7.90e^{-05}$ | .0466 |
| ENSDARG00000005841 | tnni2a.2 | 15.41668 | -.00076 | .019671 | $6.35e^{-05}$ | .0417 |
| **FBMB: SW – BW (upregulated)** | | | | | | |
| ENSDARG00000086615 | CR846087.1 | 24.25384 | 1.80942 | .584330 | $6.62e^{-05}$ | .0357 |
| ENSDARG00000104561 | znf1081 | 21.90228 | 1.89583 | .645243 | 0.00011 | .0481 |

| | | | | | | |
|---|---|---|---|---|---|---|
| ENSDARG00000106267 | CR388042.1 | 35.03077 | 5.82921 | .909595 | $9.21e^{-12}$ | < .001 |
| ENSDARG00000003989 | crhr1 | 326.8612 | 0.77404 | .245134 | $5.24e^{-05}$ | .0324 |
| ENSDARG00000096243 | ighv1-3 | 5.821612 | $8.89e^{-05}$ | .016350 | $1.61e^{-08}$ | < .001 |
| ENSDARG00000069940 | ppap2d | 846.4461 | .38736 | .113100 | $2.21e^{-05}$ | .0205 |
| ENSDARG00000044341 | chst7 | 1111.210 | .35456 | .111592 | $5.22e^{-05}$ | .0324 |
| ENSDARG00000018077 | rbp1.1 | 114.7749 | 1.77637 | .392526 | $2.40e^{-07}$ | .0006 |
| ENSDARG00000062661 | abca4b | 19.92965 | 2.46443 | .801192 | $6.43e^{-05}$ | .0357 |
| ENSDARG00000001913 | palmda | 74.66797 | 4.98298 | 1.639879 | $4.38e^{-05}$ | .0301 |
| ENSDARG00000044501 | viml | 313.7329 | 1.87359 | .447091 | $1.03e^{-06}$ | .0017 |
| ENSDARG00000037256 | si:ch211-145b13.5 | 168.8158 | .00327 | .016823 | $3.41e^{-05}$ | .0257 |
| ENSDARG00000012574 | slkb | 394.8304 | .45717 | .147247 | $6.54e^{-05}$ | .0357 |
| ENSDARG00000102888 | gpr39 | 12.16084 | 4.91526 | 1.060189 | $4.24e^{-07}$ | .0010 |
| ENSDARG00000075685 | tlr7 | 563.9174 | .62366 | .158011 | $2.88e^{-06}$ | .0043 |
| ENSDARG00000076994 | adgra2 | 29.16110 | .89893 | .307523 | 0.00012 | .0489 |
| ENSDARG00000117011 | ewsr1b | 329.9797 | 1.43001 | .454609 | $5.39e^{-05}$ | .0324 |
| ENSDARG00000025504 | gucy2f | 9.359947 | 2.64989 | .891730 | $9.61e^{-05}$ | .0454 |
| ENSDARG00000093931 | rflnb | 350.3003 | .76524 | .142432 | $3.32e^{-09}$ | < .001 |
| ENSDARG00000102356 | scp2b | 703.4846 | .29361 | .098682 | 0.00011 | .0481 |
| ENSDARG00000102050 | MCOLN3 | 22.21918 | 3.32804 | .853902 | $3.43e^{-06}$ | .0046 |
| ENSDARG00000020811 | efemp2b | 225.5847 | .89730 | .257761 | $1.75e^{-05}$ | .0174 |
| ENSDARG00000086746 | prodha | 1844.124 | .56678 | .159797 | $1.58e^{-05}$ | .0173 |
| ENSDARG00000074808 | megf6b | 45.56442 | 1.59668 | .473315 | $2.56e^{-05}$ | .0228 |
| ENSDARG00000094550 | BX649490.2 | 29.53691 | $6.00e^{-05}$ | .016350 | $4.74e^{-06}$ | .0060 |
| ENSDARG00000111856 | BX649490.4 | 27.61877 | $5.90e^{-05}$ | .016350 | $5.42e^{-06}$ | .0065 |
| ENSDARG00000045827 | lyrm5b | 22.12814 | 4.42498 | 1.394725 | $3.94e^{-05}$ | .0279 |
| ENSDARG00000002347 | cyp11a1 | 11.00258 | $6.68e^{-05}$ | .016350 | $1.54e^{-07}$ | .0005 |
| ENSDARG00000078707 | sema7a | 232.7492 | .407375 | .121829 | $2.96e^{-05}$ | .0241 |
| **FBMB: SW – BW (downregulated)** | | | | | | |
| ENSDARG00000054597 | cnot6l | 376.0056 | -.00472 | .017345 | $6.67e^{-05}$ | .0357 |
| ENSDARG00000046090 | dhrs11a | 138.8515 | -.00570 | .017810 | $8.70e^{-05}$ | .0437 |
| ENSDARG00000088753 | cfap299 | 17.16989 | -.26277 | .293151 | $7.10e^{-07}$ | .0013 |
| ENSDARG00000037805 | lgals3bpa | 55.73727 | -.00140 | .016437 | $1.32e^{-05}$ | .0152 |
| ENSDARG00000113977 | fthl29 | 15.98906 | -.00093 | .016388 | $3.00e^{-05}$ | .0241 |
| ENSDARG00000101322 | tfr1a | 258.4248 | -.00363 | .016938 | $1.81e^{-05}$ | .0174 |
| ENSDARG00000029259 | zgc:136493 | 213.7953 | -.36098 | .270908 | $2.36e^{-08}$ | < .001 |
| ENSDARG00000020084 | tg | 28.72713 | -.00133 | .016430 | $1.04e^{-06}$ | .0017 |
| ENSDARG00000020114 | slc20a1a | 306.0312 | -.00140 | .016436 | 0.00011 | .0481 |
| ENSDARG00000100020 | pim1 | 1117.054 | -.33719 | .104617 | $5.11e^{-05}$ | .0324 |
| ENSDARG00000007693 | nfkbiab | 1082.677 | -.00662 | .018379 | $3.14e^{-06}$ | .0045 |
| ENSDARG00000057644 | adam8b | 102.6395 | -.00465 | .017314 | $7.99e^{-05}$ | .0415 |
| ENSDARG00000057206 | nmt1b | 95.62002 | -.06368 | .357218 | $9.36e^{-05}$ | .0451 |
| ENSDARG00000086762 | gsdmea | 84.33024 | -.00326 | .016821 | $3.87e^{-05}$ | .0279 |
| ENSDARG00000037551 | pm20d1.1 | 38.18667 | -.00075 | .016375 | $6.82e^{-07}$ | .0013 |
| ENSDARG00000095962 | fhdc5 | 46.44994 | -.00234 | .016591 | $9.31e^{-05}$ | .0451 |
| ENSDARG00000054319 | oxct1b | 16.58725 | -.00093 | .016389 | $2.84e^{-05}$ | .024 |
| ENSDARG00000087633 | si:dkey-11o18.5 | 118.2286 | -.00080 | .016379 | $8.08e^{-09}$ | < .001 |
| ENSDARG00000060390 | stk26 | 207.8554 | -.69370 | .208613 | $3.10e^{-05}$ | .0241 |
| ENSDARG00000105511 | BX248521.2 | 38.42651 | -.00377 | .016996 | $5.26e^{-07}$ | .0012 |
| ENSDARG00000024295 | slc11a2 | 241.8629 | -.41154 | .140339 | 0.00011 | .0481 |
| ENSDARG00000104047 | si:ch211-262i1.3 | 205.5569 | -.00324 | .016816 | 0.00011 | .0481 |
| ENSDARG00000002494 | itgb6 | 42.03611 | -.00123 | .016418 | $4.01e^{-09}$ | < .001 |
| ENSDARG00000091762 | zbtb40 | 95.17587 | -.00683 | .018480 | $8.10e^{-05}$ | .0415 |
| ENSDARG00000087186 | si:ch211-232b12.5 | 899.7481 | -.55184 | .176480 | $4.72e^{-05}$ | .0316 |
| ENSDARG00000013477 | gata1a | 121.8890 | -.00332 | .016840 | $5.58e^{-05}$ | .0328 |
| ENSDARG00000026098 | si:dkey-13a21.4 | 15.25367 | -.00125 | .016419 | 0.00012 | .0489 |
| ENSDARG00000022372 | kng1 | 34.04275 | -.95705 | .598320 | $1.48e^{-12}$ | < .001 |
| ENSDARG00000101362 | mibp | 1026.365 | -.00198 | .016524 | $1.79e^{-05}$ | .0174 |
